# Multi-Omics Analysis of Fibroblasts from the Invasive Tumor Edge Reveals that Tumor-Stroma Crosstalk Induces O-glycosylation of the CDK4-pRB Axis

**DOI:** 10.1101/2021.05.28.446229

**Authors:** Gina Bouchard, Fernando Jose Garcia Marques, Loukia Georgiou Karacosta, Weiruo Zhang, Abel Bermudez, Nicholas McIlvain Riley, Lindsey Catherine Mehl, Jalen Anthony Benson, Joseph B Shrager, Carolyn Ruth Bertozzi, Sharon Pitteri, Amato J Giaccia, Sylvia Katina Plevritis

**Affiliations:** Department of Biomedical Data Science, Stanford University, Stanford, CA 94305, USA; Department of Radiology, Canary Center for Cancer Early Detection, Palo Alto CA, 94304, USA; Department of Radiation Oncology, Stanford, CA 94305, USA; Departments of Chemistry, Stanford University, Stanford, CA 94305, USA; Department of Cardiothoracic Surgery, Stanford University, Stanford, CA 94305, USA; Department of Oncology, University of Oxford, Oxford OX3 7DQ, UK

**Keywords:** Tumor-Adjacent Fibroblasts (TAFs), O-glycosylation, Tumor microenvironment (TME), multi-omics, Lung Adenocarcinoma (LUAD)

## Abstract

The invasive leading edge represents a potential gateway for tumor invasion. We hypothesize that crosstalk between tumor and stromal cells within the tumor microenvironment (TME) results in the activation of key biological pathways depending on their location in the tumor (edge vs core). Here, we highlight phenotypic differences between Tumor-Adjacent-Fibroblasts (TAFs) from the invasive edge and Cancer-Associated Fibroblasts (CAFs) from the tumor core, established from human lung adenocarcinomas. We use an innovative multi-omics approach that includes genomics, proteomics and, O-glycoproteomics to characterize crosstalk between TAFs and cancer cells. Our analysis shows that O-glycosylation, an essential post-translational modification resulting from sugar metabolism, alters key biological pathways including the CDK4-pRB axis in the stroma, and indirectly modulates pro-invasive features of cancer cells. In summary, aside from improving the efficacy of CDK4 inhibitors anti-cancer agents, the O-glycoproteome poses a new consideration for important biological processes involved in tumor-stroma crosstalk.

## Introduction

The leading edge of tumors represents one of the main site of access to local invasion and metastasis (1), however, the role of fibroblasts from the tumor edge have been mainly overlooked. Much of our current understanding of cancer-associated fibroblasts (CAFs) is derived from fibroblasts harvested from the inner tumor, and the pro- and anti-tumor effects of CAFs are still being resolved (2–5). Single-cell studies have demonstrated significant CAFs heterogeneity in the tumor microenvironment (TME) (6,7) but the cell-cell interactions involving different tumor fibroblast subtypes, particularly those at the tumor edge, are not well known. Of the few studies that focus on fibroblasts at the invasive tumor edge, colorectal and hepatocellular carcinomas studies found peritumor fibroblasts to stimulate the proliferation and migration of cancer cells more strongly than CAFs (8,9). In order to better understand the role of these peritumor fibroblasts in tumor invasion and treatment response, fibroblast heterogeneity and cell-cell interactions at the invasive edge of the tumor need to be further explored.

While multi-omics analysis of fibroblasts have included epigenomic, genomic and transcriptomic data (10–12), post-translational modifications (PTM) of fibroblasts have not been well characterized. PTMs have gained significant interest in recent omics studies as they directly modulate numerous cancer hallmarks including cell cycle progression, migration and metabolism (12–15). Previous work showed that metabolic reprogramming toward the hexosamine biosynthesis pathway (HBP), an understudied glucose pathway leading to glycosylation, is associated with poor survival in the lung adenocarcinoma (LUAD) (16,17). The authors reported that this metabolic switch happens earliest in CAFs compared to the malignant cells across tumor compartments (fibroblasts, cancer, immune and endothelial cells). This finding suggests that tumor fibroblasts redirect their glucose toward HBP, leading to altered O-glycosylation, a PTM regarded to be as critical as phosphorylation and also initiated on serine and threonine residues (18,19). While previous results on CAFs assessed metabolic reprogramming toward HBP in the tumor stroma, direct measurements of the O-glycoproteome changes following tumor-stroma interactions using a multi-modal approach have yet to be reported.

O-linked glycosylation has been associated with several cancer hallmarks (20,21). However, very little is known about: 1) which proteins are O-glycosylated 2) the effect(s) of O-glycosylation on protein function, and 3) how protein O-glycosylation impacts cell-cell crosstalk. To our knowledge, no published study has reported the O-glycoproteome of tumor fibroblast nor multi-omics approach that integrates the O-glycoproteome in the context of tumor-stroma crosstalk. The lack of data on the O-glycoproteome is largely due to the technical challenges associated with probing glycobiology (22) including the lack of automated techniques, small libraries of glycoproteins, and limited availability of sugar-specific antibodies (Huang et al, 2020; National Research Council, 2012). Currently, mass spectrometry is the main method used to measure O-glycosylation but, the lability of O-glycans, the lack of universal enrichment methods due to O-glycan heterogeneity, and the limitations of bioinformatic search engines for O-glycopeptides (searching time and accuracy), as well as their quantification, add to the technical challenges. However, recent enrichment methods (24,25), employment of O-glycoproteases (26–28) and the SimpleCell approach that prevents O-glycan elongation and offers a simplified version of the O-glycoproteome (29) have allowed us to improve the characterization and investigate the role of O-glycosylation in the TME.

This study focuses on the understudied peritumor fibroblasts from the invasive edge of primary LUAD tumors referred hereon as Tumor-Adjacent-Fibroblasts (TAFs) and demonstrates the unconventional phenotype of TAFs compared to CAFs harvested from the inner tumor. We investigate the role of TAFs through multi-omics profiling of tumor-stroma cocultures by integrating transcriptomic, proteomic, and for the first time, O-glycoproteomic data obtained by an optimized method of O-glycan enrichment (25) and precise spectra quantification of O-glycopeptides (30). We find that the O-glycoproteome provides additional biological insights on tumor-stoma crosstalk over genomic and proteomic data and highlights the role of O-glycosylation in modulating the CDK4-pRB axis of fibroblasts at the tumor invasive edge, among other important biological processes. Lastly, we show that targeting O-glycosylation in TAFs indirectly decreases pro-invasive features in cancer cells, establishing the O-glycoproteome as an important way to modulate the TME.

## Results

### The fibroblasts from the invasive tumor edge have a distinct phenotype compared to those from tumor core

Fibroblasts from the leading edge of tumors have been shown to facilitate invasion of cancer cells (9) but these fibroblasts have not been fully characterized. To better understand fibroblasts from the invasive tumor edge, we established matched primary fibroblast cell lines from different tumor regions namely Normal Fibroblasts (NFs, >5 cm away from the tumor), Tumor-Adjacent Fibroblasts (TAFs, invasive tumor edge) and Cancer-Associated Fibroblasts (CAFs, tumor core) (Fig. 1A) from fresh human LUAD clinical specimens (Fig. S1A for clinical annotations). We find that the three subtypes of fibroblasts present striking morphological differences (Fig. 1B). NFs demonstrate the typical spindle-like morphology commonly associated with fibroblasts and CAFs show enlarged cellular bodies, which agree with the commonly described “activated” fibroblast morphology. In contrast, TAFs are filiform with a striated organization, a distinct morphology for tumor fibroblasts. We then profiled the NFs, TAFs and CAFs using flow cytometry and RNA sequencing (RNA-seq). To visualize the flow cytometry data, we applied the high-dimensional embedding tSNE algorithm to project α-SMA, FSP1, GFAT2, CD10, p21, and vimentin markers for each cell onto a two-dimensional space. NFs, TAFs and CAFs clustered into distinct regions on the tSNE plot with some overlap between NFs and TAFs (Fig. 1C). For each clinical specimen, α-SMA and FSP1 were consistently high in CAFs, intermediate in the NFs and low in TAFs; the other markers were more variable across specimens (Fig. S1B and S1C). From the RNA-seq data, we found that none of the canonical CAF markers were significantly differentially expressed among the different fibroblast subtypes (Fig. S1D and S1E) but the functional gene set enrichment analysis revealed that NFs were uniquely enriched for glycogen degradation; TAFs, for the thioredoxin redox metabolism, as well as lipid signaling mediated by Tubby pathways; and CAFs, for inflammatory and pro-fibrotic pathways including IL-10 and NF-κB signaling (Fig. 1D, E and F). Of note, all fibroblast subtypes, including NFs, were enriched for pathways associated with a stress response (p38 MAPK), suggesting that our NFs may also be partially transformed (Fig. 1F).

**Figure 1.**
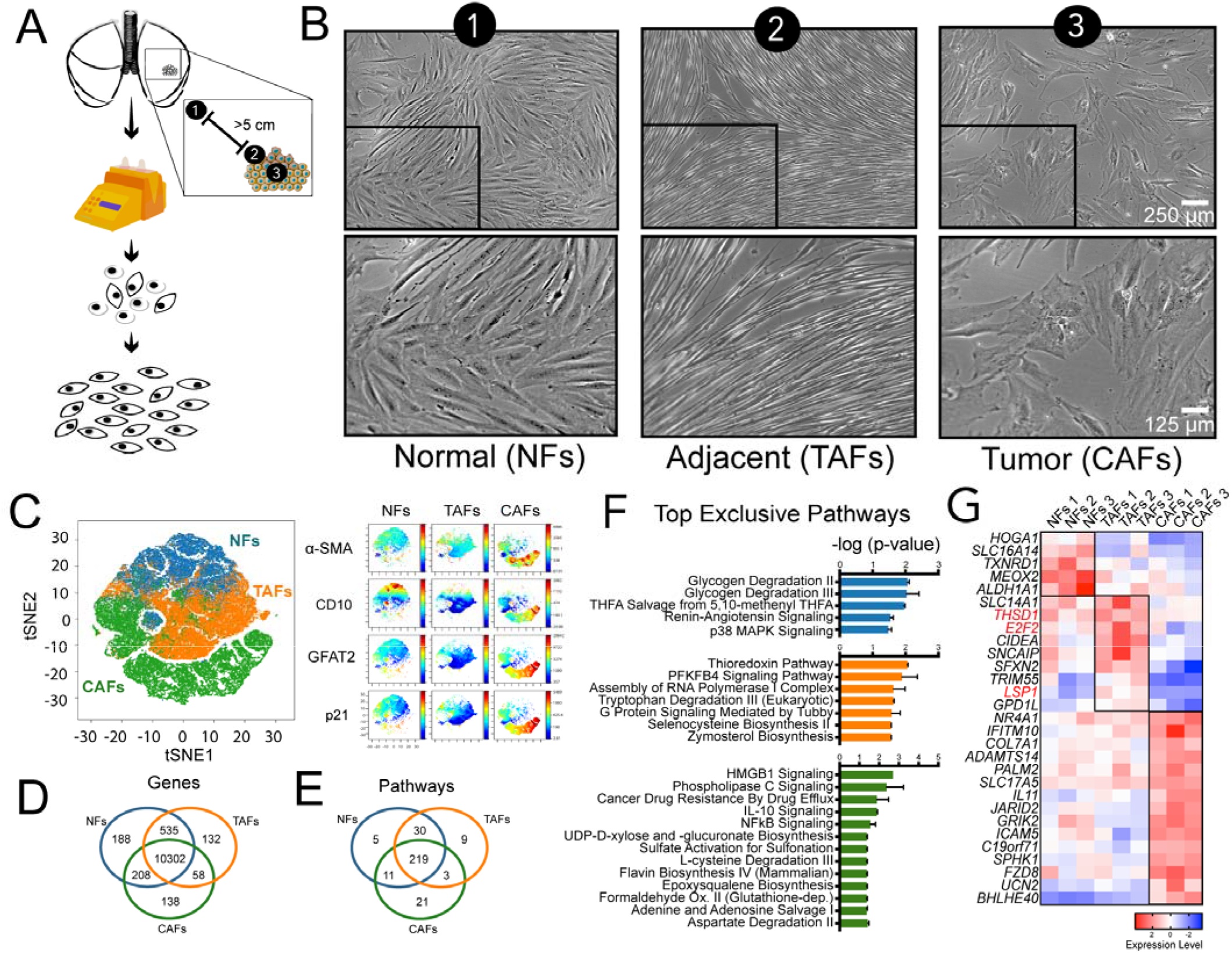
Tumor region analysis of LUAD fibroblasts. **A**, Fibroblast expansion workflow of LUAD fresh specimens. **B,** Phase contrast microscopy images of NFs, TAFs, and CAFs. **C**, Representative viSNE analysis of NFs, TAFs, and CAFs using with α-SMA, FSP1, GFAT2, CD10, p21, and vimentin as clustering markers. Other biological replicates available in Fig. S1C. Venn diagram showing the gene overlap measured by RNAseq (TPM >1 in all biological replicates of each subtype) (**D)** Venn diagram showing pathway overlap (**E**) and bar graph (**F**) showing the top exclusive pathways in NFs (n=3), TAFs (n=3), and CAFs (n=3) analyzed with Ingenuity Pathway Analysis software. **G**, Heatmap showing relative normalized gene expression of top DEGs in NFs, TAFs, and CAFs.

Several interesting markers were identified among the genes most uniquely expressed in TAFs, as compared to NFs and CAFs, including the E2F2 transcription factor (Fig. 1G), known for its role in proliferation and association with a poor prognosis in LUAD (31,32). TAFs also overexpressed Thrombospondin Type 1 Domain Containing 1 (THSD1), a marker of hematopoietic stem cell (33) and the fibrocyte marker leukocyte-specific protein 1 (LSP1). In addition, TAFs show a moderate expression of α-SMA (Fig. 1C, S1C, S1D and S1E) as well as a significant increase in antigen presentation and lipid signaling compared to CAFs (Fig. 1F and S1G), which are also known fibrocyte properties (34). This fibrocyte signature is intriguing because fibrocytes are antigen-presenting leukocytes derived from peripheral blood mononuclear cells implicated in wound healing and displaying fibroblast-like properties. Fibrocytes are present in injured organs and have both inflammatory features of macrophages and tissue remodeling properties of fibroblasts (34). Taken together, these results show a distinct subtype of tumor fibroblasts, namely TAFs, which display some fibrocyte-like properties and suggest a hematopoietic origin.

### TAFs, but not CAFs, induce a mesenchymal phenotype in HCC827 LUAD cell line

To understand how the fibroblasts isolated from different tumor regions affect cancer cells, we analyzed lung cancer adenocarcinoma cells, using the HCC827 cell line, after cell-cell contact cocultures for 24 hours with NFs, TAFs and CAFs, and performed RNAseq analysis on the sorted compartments (Fig. 2A). HCC827 cells were chosen for their well characterized epithelial-to-mesenchymal (EMT) transition under TGF-β stimulation at the single-cell resolution using mass cytometry (35). Strikingly, the images of HCC827 cocultured with TAFs and CAFs greatly differ: HCC227 intermixed with TAFs, but HCC827 formed tightly packed islands of epithelial cells when cocultured with CAFs (Fig. 2B and S2A). Then, we performed gene set enrichment analysis on the differentially expressed genes (DEGs) obtained from the sorted cancer cells cocultures relative to their matched monoculture and highlighted the differences in functional enrichment between HCC827 cocultures with TAFs versus CAFs (Fig. 2C-E). Of note, cancer cells cocultured with the TAFs showed a significant increase for pathways associated with migration, invasion, and micropinocytosis, among others; in contrast, CAFs promoted functional enrichment of differentiation pathways and glycosaminoglycan production in the cancer cells. These transcriptional differences are consistent with the morphological differences observed in the HCC827s in the cocultures with TAFs vs CAFs.

**Figure 2.**
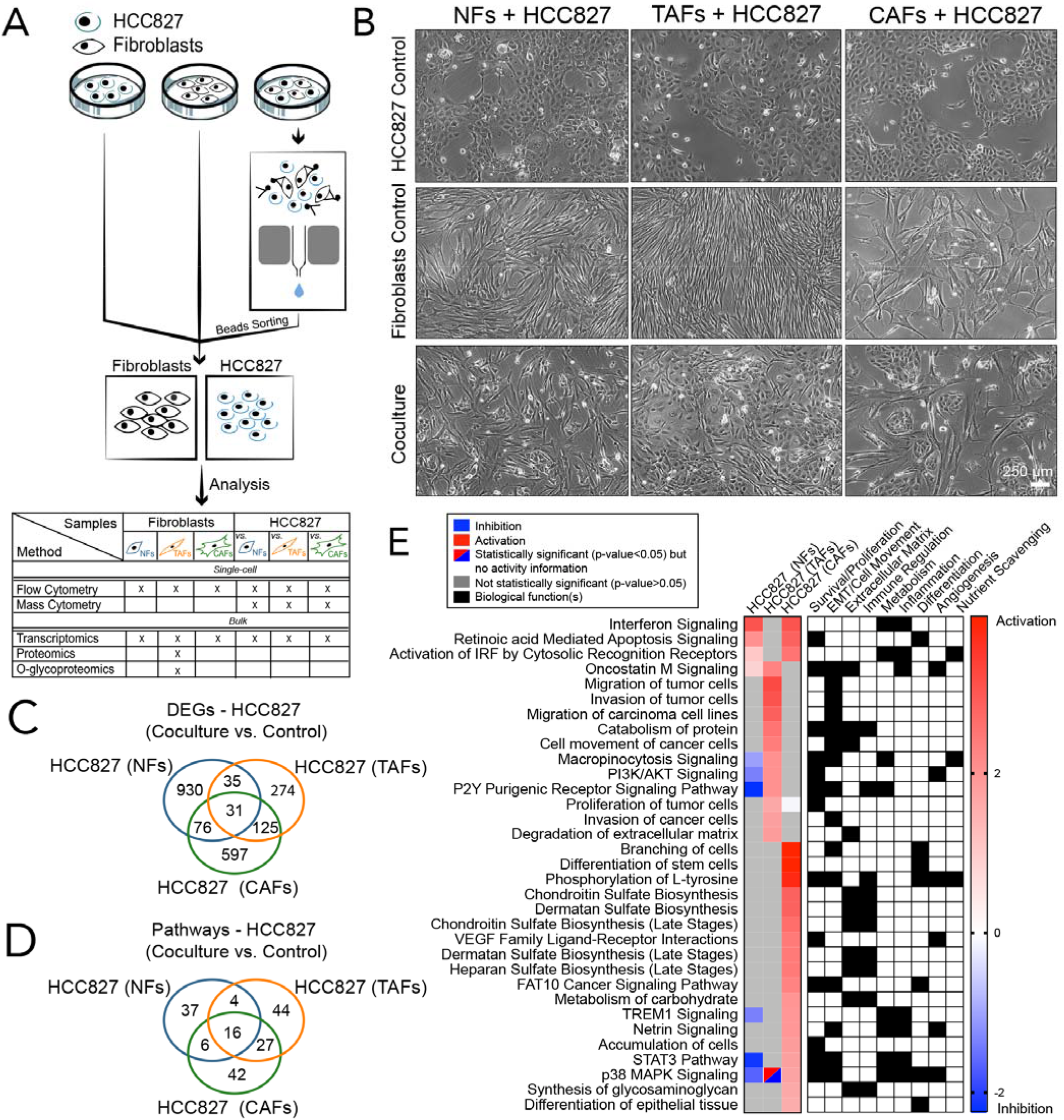
RNAseq analysis of HCC827 after coculture with NFs, TAFs and CAFs. **A**, Coculture, cell separation and downstream analysis workflow. **B**, Phase contrast microscopy images of NFs, TAFs and CAFs cocultures with HCC827 lung adenocarcinoma cell line. **C**, Venn diagram showing the gene overlap between HCC827 cocultures with NFs (n=3), TAFs (n=3) and CAFs (n=3) compared to their monoculture control measured by RNAseq (p < 0.05, fold-change > 0.5). Venn diagram showing the pathways overlap and (**D**), heatmap showing the main pathways/biological functions differences between HCC827 cocultures with NFs (n=3), TAFs (n=3) and CAFs (n=3) compared to their monoculture control (**E**) analyzed with the IPA software.

Next, HCC827 cocultures were analyzed with mass cytometry (CyTOF) using an EMT panel of markers and projected on PHENOSTAMP, a publicly available lung cancer reference map of EMT phenotypic states (35). Only the TAFs significantly induced EMT in the HCC827, as illustrated on the PHENOSTAMP projections (Fig. 3A and B). In addition, we used the forced-directed layout and X-shift clustering algorithm to visualize the mesenchymal cell populations after coculture (Fig. 3C and S2B-S2D). Again, only the HCC827 after direct cell-cell contact coculture with TAFs showed a mesenchymal phenotype, as confirmed by the position of the HCC827 cocultured cells (dark green cluster) with the other mesenchymal cells (Fig. 3C). HCC827 showed a significant decrease of epithelial markers, transcription factors, and signaling, as well as an increase of mesenchymal markers after direct contact with TAFs. Interestingly, supernatant from TAFs did not promote a significant EMT response in HCC827, suggesting that direct contact between cancer cells and TAFs is necessary to induce the mesenchymal phenotype in the cancer cells (Fig. S3A). In summary, our results suggest that TAFs induce a greater mesenchymal phenotype in cancer cells compared to NFs and CAFs, which is mediated through direct cell-cell contact.

**Figure 3.**
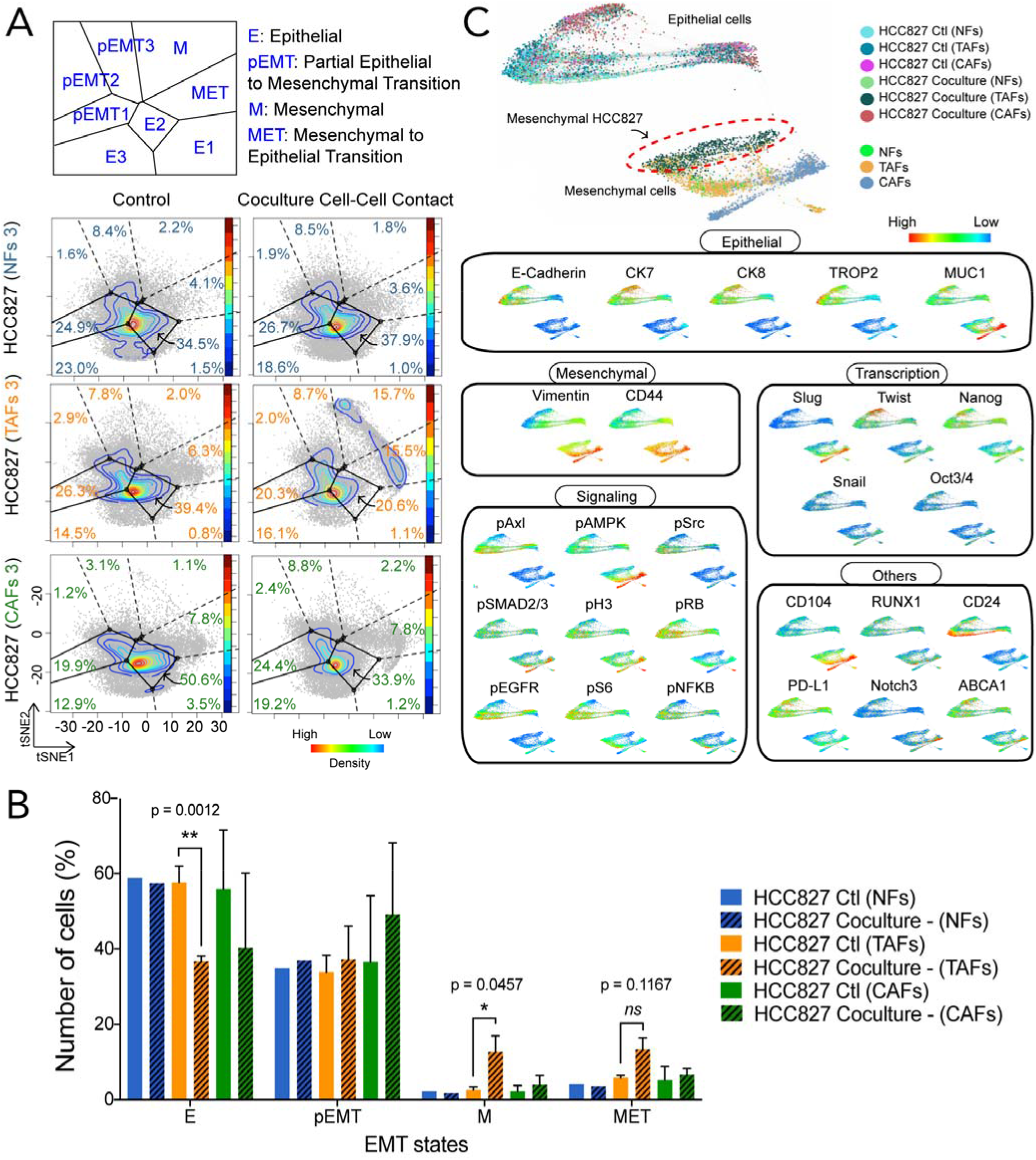
CyTOF analysis of HCC827 after coculture with NFs, TAFs and CAFs. Projections of the HCC827 mococulture controls and matched cocultures after cell-cell contact with NFs, TAFs and CAFs onto the EMT–MET PHENOSTAMP phenotypic map (**A**) and bar graph quantification (**B**) of the number of cells in each region of the map. **C**, Single-cell force-directed layout colored by protein expression of indicated markers of HCC827 cocultures with NFs, TAFs and CAFs using X-shift clustering (arcsinh transformed data of specimen 3). See Fig. S2B for expression scale bars and S2C and S2D for biological replicates.

### HCC827 cells induce differential metabolic reprogramming in TAFs vs CAFs

To measure the effect of cancer cells on primary fibroblasts, we then analyzed the transcriptome of NFs, TAFs, and CAFs 24 hours after coculture with HCC827 cells (Fig. 4A, B and S3B-S3D). Gene set enrichment analysis shows that TAFs experience a significant decrease in cell movement and signaling, particularly lipid signaling (inositol compounds). Conversely, CAFs experience an increase in most of these pathways, as well as pathways associated with metabolism and immune regulation (Fig. 4C). Overall, TAFs activity is drastically downregulated without significantly affecting cell death (Fig. S3E), as opposed to almost all of the same pathways significantly upregulated in NFs and CAFs cocultures (Fig. S3B).

**Figure 4.**
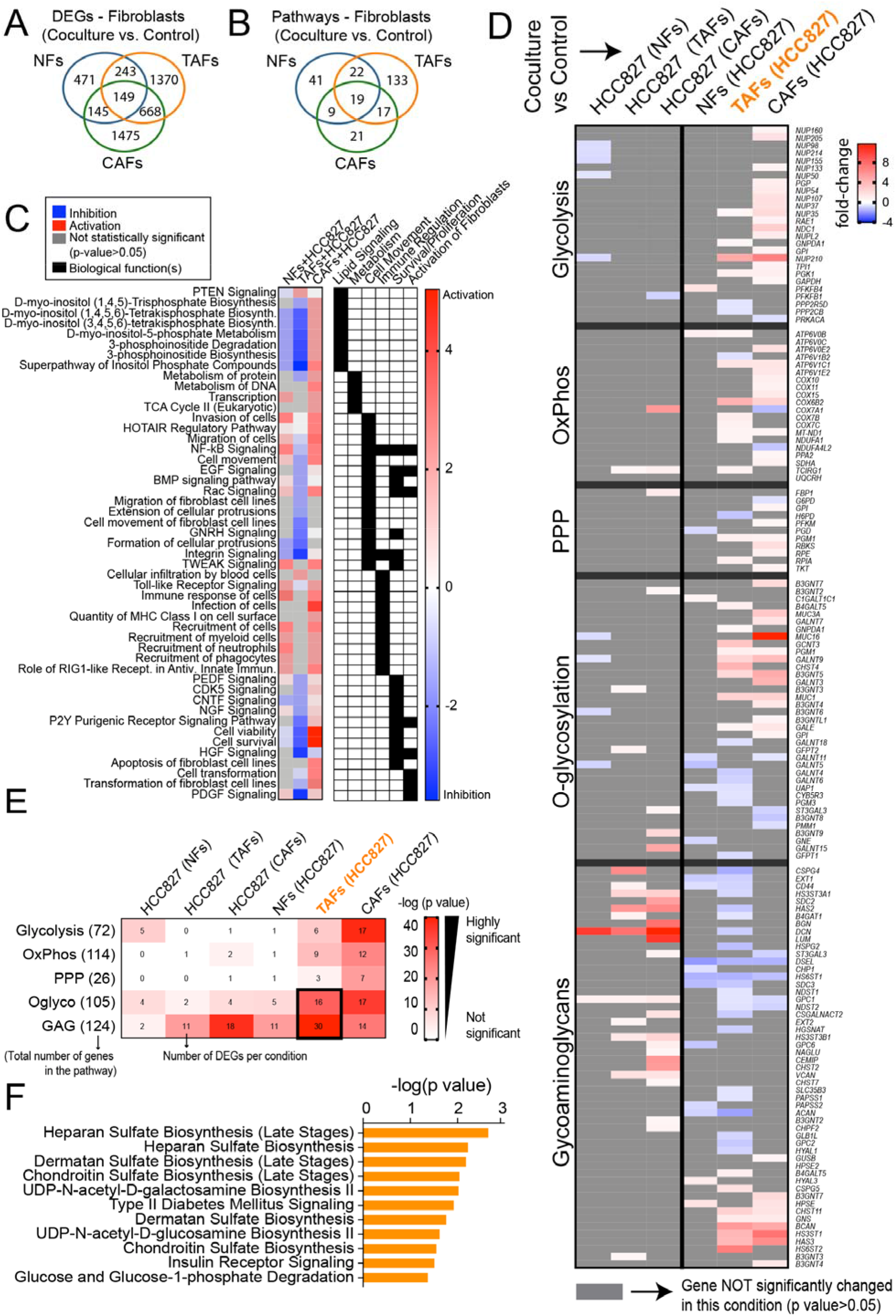
Gene expression analysis of NFs, TAFs and CAFs after coculture with HCC827 lung adenocarcinoma cancer cells. **A**, Venn diagram showing the gene overlap between NFs (n=3), TAFs (n=3) and CAFs (n=3) cocultures with HCC827 compared to their monoculture control measured by RNAseq analysis (p < 0.05, fold-change > 0.5). Venn diagram showing pathway overlap (**B**), heatmap showing the main pathways/biological functions differences analyzed with the IPA software (**C**) and, heatmap showing the fold-change of DEGs in the main glucose metabolic pathways between NFs (n=3), TAFs (n=3) and CAFs (n=3) cocultures with HCC827 (**D**) and heatmap showcasing the number of overlap genes between DEGs in each condition within each metabolic pathway (**E**). Color of heatmap represents -log(p-value) of the gene overlap calculated using a hypergeometric test. The number in parentheses show the total number of genes from each MSigDB gene set and tile numbers show the number of DEGs for each condition. See p values in Fig. S3F. **F,** Bar graph showing exclusive pathways related to glycans biosynthesis enriched in TAFs cocultures compared to their monoculture control and analyzed with the IPA software.

Prior work from our group reported transcriptional alterations in CAFs and metabolic reprogramming toward HBP in primary LUAD tumors (16). Here, we explored the significant DEGs involved in metabolic reprogramming for all cocultures in the main glucose pathways (glycolysis, oxidative phosphorylation, pentose phosphate pathway, O-glycosylation and glycosaminoglycans biosynthesis) using the Molecular Signatures Database (MSigDB) gene sets. Strikingly, transcriptomic changes in metabolic pathways occurs primarily in tumor fibroblasts as opposed to cancer cells (Fig. 4D, E and S3F), as illustrated by the low number of genes significantly changed in cancer cell cocultures (relative to monoculture control). As expected from previously published work, several metabolic pathways were found to be significantly dysregulated in CAFs (Fig.4E) but interestingly, we show that the main pathways modulated in TAFs were related to glycan biosynthesis. We also highlight several other exclusive pathways related to glucose metabolism with unknown activity in the TAFs cocultures, including glucose degradation, nucleotide sugars and glycoproteins biosynthesis, all of which were identified through the Ingenuity Pathway Analysis (IPA) software (Qiagen, Fig. 4F). These transcriptomic results suggest that fibroblasts from the invasive edge of the tumor undergo differential metabolic reprogramming compared to CAFs. In particular, TAFs upregulate glycoproteins biosynthesis when in direct cell-cell contact with cancer cells, suggesting an important role of glycans at the tumor invasive edge.

### The TAFs O-glycoproteome reveals previously unidentified O-glycoproteins

Our transcriptomic analysis suggests that the hexosamine (aminosugars biosynthesis leading to O-glycosylation) and glycan biosynthesis pathways in TAFs are modulated by tumor-stroma crosstalk. Because these phenomena are largely uncharacterized, we assessed the effects attributed to cancer cells-TAFs crosstalk through mono- and coculture analysis on the proteome and O-glycoproteome using mass spectrometry. Briefly, TAFs were sorted after coculture with HCC827, then their protein extracts were digested with trypsin and enriched for O-glycopeptides using an optimized method based on previously published work (25).

For a control, a matched shotgun proteomic sample not subjected to O-glycopeptide enrichment was prepared and analyzed for each sample Across all the conditions, the proteomic analysis revealed 4495 unique proteins in which 337 O-glycoproteins were found to be O-glycosylated on 780 putative O-glycosites (Fig. 5A). Of the 337 O-glycoproteins identified in TAFs, 43 proteins were found to be differentially O-glycosylated (adjusted for protein level differences) after coculture with HCC827, most of them unknown O-glycoproteins according to the most comprehensive repository, the OGP database (23). The majority of these proteins were enzymes, transcription regulators, transporters or cytoskeleton proteins (Fig. 5B). O-glycans are classified into common core structures depending on their composition and sugar configuration (Tn antigen and core 1 to 8). Of the 43 differentially O-glycosylated proteins measured here, core 1 O-glycans, the most common core structure (36), were the most abundant (Fig. 5C). In addition, approximatively 50% of the O-glycans were sialylated, a common glycan modification in cancer that blocks further linear extension of O-glycan chains (36). Overall, we generated the first stromal O-glycoproteome in the context of tumor-stroma crosstalk with their putative O-glycosites, providing a new resource for developing targeted O-glycosylation strategies and tools.

**Figure 5.**
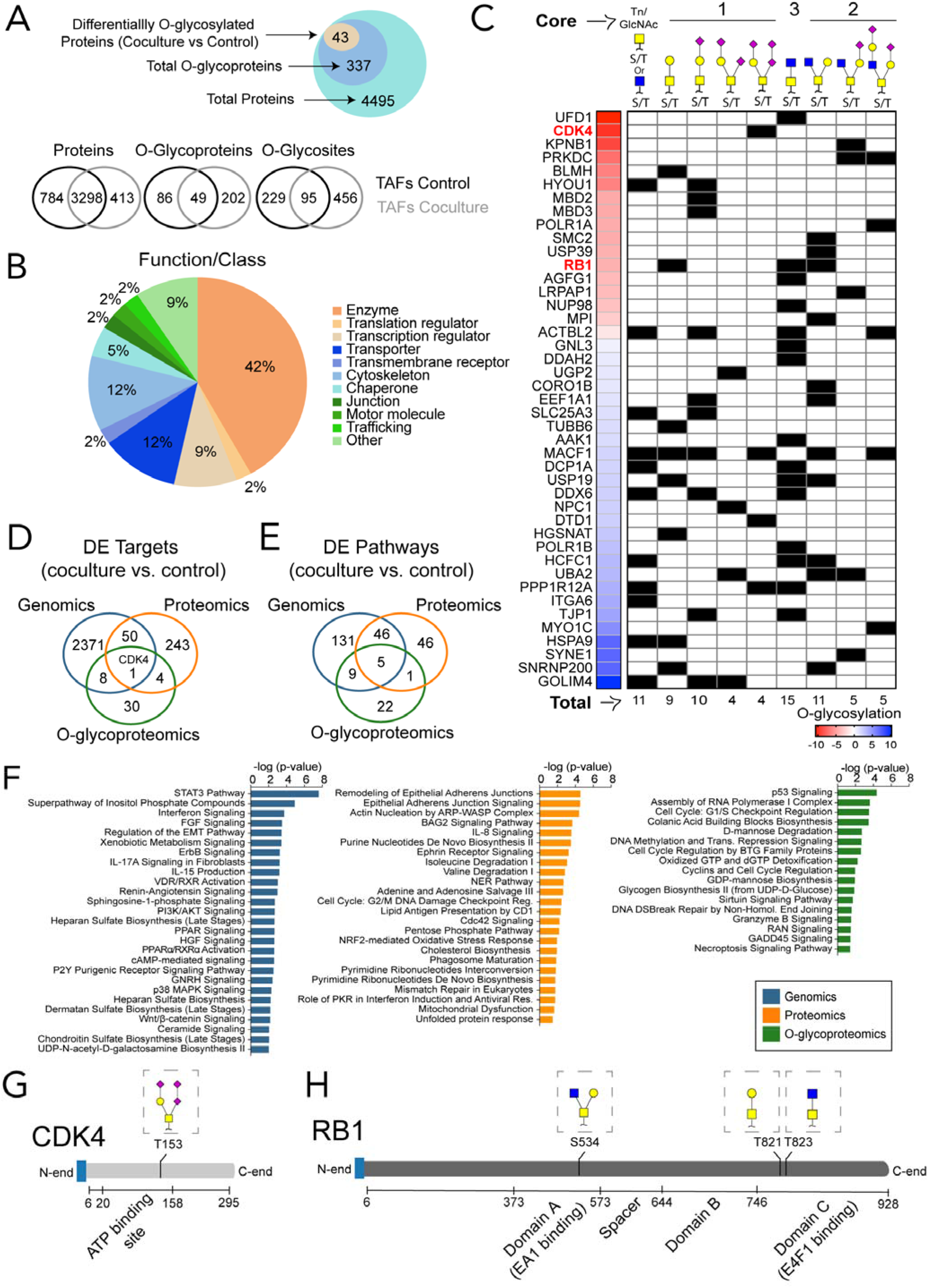
Crosstalk with HCC827 disrupts the TAFs O-glycoproteome. **A**, Venn diagram showing proteins, O-glycoproteins, and O-glycosites found in TAFs monocultures and cocultures analyzed by mass spectrometry. **B**, Pie charts showing the main classes of the differentially O-glycosylated proteins after coculture identified by mass spectrometry. **C**, Heat map and summary table of the differentially O-glycosylated proteins and glycan heterogeneity after coculture with HCC827. Venn Diagram showing the overlap of targets (**D**) and the DE pathways identified in TAFs cocultures (relative to control monoculture) (**E**) using different methods of analysis. **F,** Bar graph highlighting a sample of the top enriched pathways in TAFs cocultures at the genomic, proteomic, and O-glycoproteomic level. CDK4 (**G**) and RB1 (**H**) protein domains and O-glycan positions.

Next, the differentially expressed proteins and O-glycoproteins were analyzed for pathway enrichment and compared with the transcriptomic data (Fig. 5D-F). The pathways measured with different assays show a modest overlap suggesting that each data dimension highlights unique molecular functional characterization in TAFs cocultures. Interestingly, the activity measured for shared pathways did not necessarily increase or decrease conjointly at the gene and protein level, highlighting the relevance of a multi-omics approach to more accurately interpret biological activity (Fig. S3G). Then, we compared the top unique pathways (Fig. 5F) and summarized the biological functions for each method of analysis. Briefly, cell proliferation, immune response, and cell movement are regulated primarily at the gene expression level; energy, amino acids and nucleotide metabolism as well as cytoskeleton remodeling are regulated at the protein level; and lastly, O-glycosylation regulates cell cycle, cell death and glucose, mannose and guanosine metabolism. Taken together, these results support the use of a multi-omics approach that includes the transcriptomics, proteomics and O-glycoproteomics to more fully characterize biological functions.

### O-glycosylation regulates the CDK4-pRB axis in TAFs and modulates the anti-cancer properties of CDK4/6 inhibitors

To validate the relevance of the stromal O-glycoproteome in tumor-stroma crosstalk, we explored the role of O-glycosylation in the pRB-CDK4 axis as an example since this signaling pathway stood out from our analysis. Briefly, cyclin dependent kinase 4 (CDK4) phosphorylates the tumor suppressor retinoblastoma protein 1 (RB1), leading to its inactivation and uncontrolled cell proliferation. In our multi-omics analysis, only one target, CDK4, was found to be differentially upregulated at the gene, protein and O-glycosylation level in the TAFs after coculture with HCC827 cells (Fig. 5D). In addition, both CDK4 and RB1 were found to be O-glycosylated at a higher level in TAFs after direct cell-cell contact with cancer cells. Interestingly, CDK4 is O-glycosylated at T153 in TAFs cocultures (Fig. 5G and S3H), which is expected to be in the active site of the enzyme (37). Moreover, upon coculture, RB1 is O-glycosylated at S534, T821 and T823 glycosites (Fig. 5H), which are predicted to be located in transcription factor interaction domains of the protein. Given the role of EMT in resistance to CDK4 inhibitors (38), the pro-EMT effect of TAFs on cancer cells observed in Fig. 3, and the location of the O-glycans on CDK4 and RB1, we hypothesized that O-glycosylation of the stroma could affect the efficacy of CDK4/6 inhibitors, a recent family of anti-cancer agents targeting the CDK4-pRB axis (39).

Unfortunately, no specific O-glycosylation inhibitors to these sites are available nor specific antibodies for O-glycans due to the limited knowledge and tools available for O-glycobiology. In addition, primary fibroblast mutants from human clinical specimens are notoriously challenging to generate due to their limited lifespan. To test this hypothesis, we then treated TAFs:HCC827-eGFP cocultures with palbociclib, a clinically-approved CDK4/6 inhibitor, alone and in combination with 6-diazo-5-oxo-L-norleucine (DON), a glutamine analog that inhibits HBP and reduces O-glycosylation, using drug concentrations in accordance with previous studies performed in non-small cell lung cancer cell lines (40,41). The main goal of this experiment was to validate the link between O-glycosylation and the efficacy of CDK4 inhibitors in the tumor stroma by measuring the phosphorylation of RB by CDK4 with and without O-glycosylation. To quantify the effect of palbociclib alone and in combination with DON, we measured the proliferation marker Ki67 and phosphothreonine 826, the CDK4-preferred phosphosite on RB1 (uniportorg) in the cocultures at 48 hours post-treatment at single-cell resolution. We also used the marker p21 to discriminate proliferative from senescent cells since p21 stabilizes the Cyclin D/CDK4 complex leading to RB phosphorylation but is also a marker of cell senescence (42,43). Briefly we focused our analysis on two main cell populations, referred as: (i) “proliferative” cells, defined as {Ki67^high^, pRB^high^, p21^high^}-cells, and (ii) “senescent” cells, defined as {Ki67^low^, pRB^high^, p21^high^}-cells that were identified on a forced-directed layout projection of two independent experiments (biological replicates) using TAFs clinical specimens from different patients. This method of analysis was primordial as the cell state (proliferative vs non-proliferative) and cell signaling through PTM are intimately linked, which resolution would be lost at the bulk level and would not allow proper analysis of the CDK4-pRB axis (44). The cocultures treated with palbociclib in combination with DON exhibited a decrease of proliferative TAFs and an increase of senescent TAFs (Fig. 6A-E) compared to palbociclib or DON alone. This effect was not observed in the monocultures (Fig. S4). Moreover, the treatment with DON and palbociclib led to a CAF-like morphology change in the TAFs, consistent with increased senescence and size. These results indicate that O-glycosylation of the CDK4-pRB axis modulates the effect of palbociclib in TAFs cocultured with cancer cells, which effect is significantly improved when inhibiting O-glycosylation with DON.

**Figure 6.**
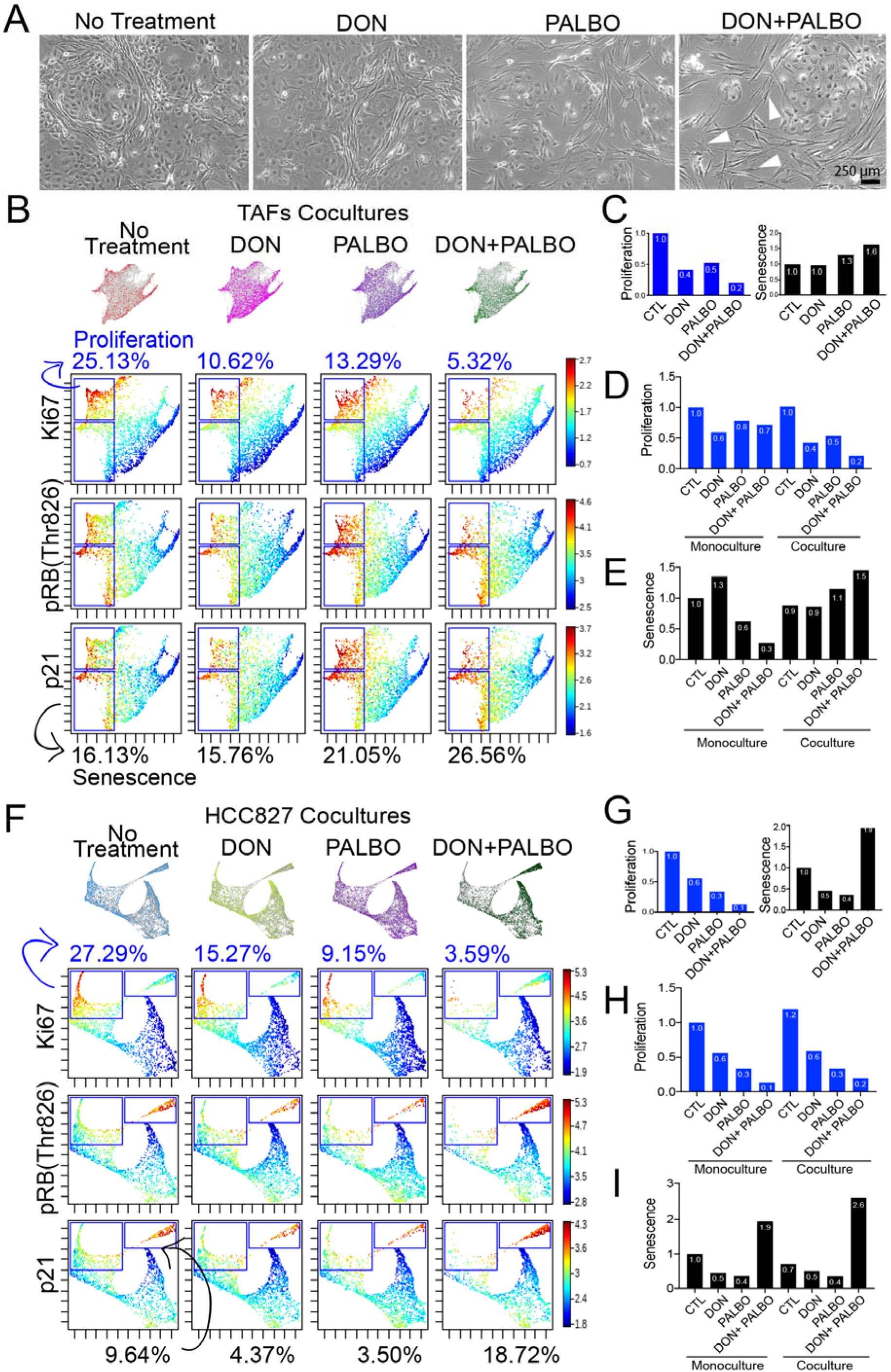
O-glycosylation of the CDK4-Rb axis in TAFs modulates Palbociclib efficacy. **A**, Phase-contrast microcopy of TAFs:HCC827 cocultures treated with 10 μM DON, 1 μM palbociclib or a combination of both. White arrowheads show TAFs that have gained an activated CAFs-like morphology. Force-directed layout of TAFs cocultures showing proliferative (upper gate) and senescent (lower gate) TAFs (**B**) and bar graph quantification of percentage ratios relative to control coculture (no treatment) **C**. Bar graph quantification showing all conditions of TAFs mono- and cocultures relative to control monoculture (no treatment) for (**D**) proliferation and (**E**) senescent TAFs. Force-directed layout of HCC827 cocultures showing proliferative (left gate) and senescent (right gate) cancer cells (**F**) and bar graph quantification of percentage ratios relative to control coculture (no treatment) **G**, Bar graph quantification showing all conditions of HCC827 mono- and cocultures relative to control (no treatment) monoculture for (**H**) proliferation and (**I**) senescent cancer cells. Results are from two independent experiments with biological replicates combined (TAFs from specimens 1 and 3).

### Modulating O-glycosylation of the CDK4-pRB axis in TAFs indirectly decreases the mesenchymal features of cancer cells

It is crucial to understand the effect of treatment in the stroma but the cancer cells remain the ultimate target in cancer. Although we did not measure the HCC827 O-glycoproteome which was not significantly altered at the transcriptomic level, we did measure proliferation, senescence and mesenchymal features in the cancer cells following the combination treatment. Briefly, our analysis suggests that the combination of palbociclib and DON is more efficacious than palbociclib alone in reducing cancer cell proliferation. However, O-glycosylation of the CDK4-pRB axis in the cancer cells does not seem to be significantly modulated by cell-cell contact with TAFs since no significant difference was observed between mono- and cocultures treated with DON (Fig. 6F-I and S5).

Lastly, to assess whether or not the TAFs morphological transformation into CAF-like cells observed in the cocultures treated with DON and palbociclib affects the cancer cells, we analyzed EMT by measuring the expression of CD44 in the cocultures by immunofluorescence (IF). Our rationale for analyzing the mesenchymal marker CD44 was based on our CyTOF analysis presented in Fig. 3, which shows a significantly lower expression of CD44 in the HCC827 cocultured with CAFs compared to TAFs and was the main mesenchymal feature associated with TAFs-induced EMT. The IF staining supports our hypothesis that TAFs may display a CAF-like phenotype with the combination treatment by showing a strong decrease of CD44 in the HCC827 when cocultured with TAFs, and by displaying a pattern resembling CAFs:HCC827 coculture morphology (Fig. 7 and S6). However, more validations and quantifications are needed to confirm the effect of the treatment alone versus the indirect effect of the treated TAFs on the cancer cells. Overall, these results suggest that not only inhibiting O-glycosylation in combination with palbociclib reduces pro-tumor properties in TAFs and HCC827, but also that these changes in the stroma are likely to decrease the mesenchymal features in cancer cells.

**Figure 7.**
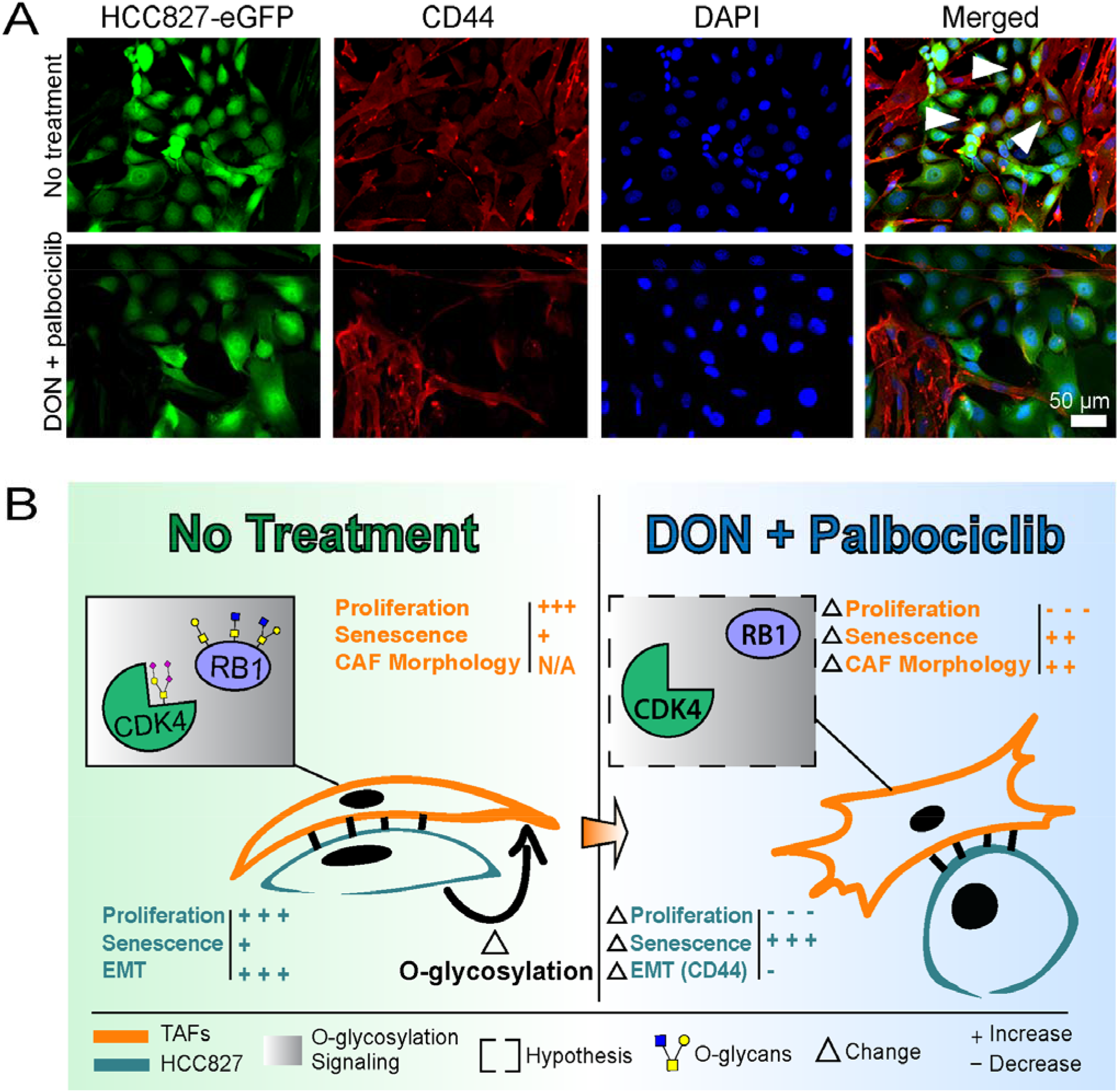
Decreased O-glycosylation with DON treatment indirectly prevents EMT in HCC827. **(A**) Immunofluorescence staining of HCC827-eGFP:TAFs coculture showing a decrease of the mesenchymal marker CD44 in cancer cells after treatment with DON and palbociclib. Cancer cells are represented by green pseudo color. TAFs are represented by red pseudo color only. CD44+ cancer cells are represented by co-staining with green and red pseudo colors. White arrowheads represent examples of CD44+ cancer cells. (**B**) Summary figure of the main findings of the study showing that EMT in HCC827 and O-glycosylation of the CDK4-pRB axis in the stroma are induced in TAFs:HCC827 cocultures, which is prevented with the combination treatment with DON and palbociclib.

## Discussion

The role of tumor fibroblasts at the tumor leading edge have been understudied. Moreover, studying fibroblasts in general is challenging for several reasons, including their phenotypic plasticity and lack of adequate markers. Here we have characterized tumor-adjacent fibroblasts (TAFs) with a multi-omics approach that includes PTMs through O-glycoproteomics. We use single-cell mass cytometry to demonstrate the increased ability of TAFs to induce EMT in cancer cells compared to CAFs, as well as the pro-migratory role of the stroma at the tumor leading edge. In addition, our transcriptomic analysis highlights that metabolic reprogramming at the leading edge is happening mainly in the stroma as opposed to cancer cells, and can be measured in the context of tumor-stroma crosstalk through the O-glycoproteome. Our multi-omics analysis of TAFs highlights O-glycosylation of the CDK4-RB axis as a consequence of tumor-stroma crosstalk and suggests a new avenue for improving the efficacy of CDK4/6 inhibitors on cancer cells by modulating TAFs. Lastly, our study generated the first stromal O-glycoproteome of more than 300 O-glycoproteins in the context of tumor-stroma crosstalk with their putative O-glycosites, providing a new resource for developing targeted O-glycosylation strategies and tools.

Compared to TAFs, CAFs have been extensively studied. Here we highlight key differences between CAFs and TAFs, derived from primary LUAD. Our transcriptomic analysis showed an increase of invasion and cell movement signatures in HCC827 when cocultured with TAFs vs CAFs. Our CyTOF experiment confirmed the ability of TAFs (and not CAFs) to induce EMT more robustly in cancer cells in coculture experiments, which was only observed with direct cell-cell contact between TAFs and HCC827. This result was compelling since TAFs express low to undetectable levels of the typical CAFs markers, but is consistent with a recent murine study. Zhao and al recently showed that TAFs, which they refer to as “peri-tumor fibroblasts,” isolated from human hepatocellular carcinoma specimens promoted proliferation and migration of cancer cells at a greater level compared to CAFs in a cancer mouse model (8). This effect was driven by STAT3 signaling, which we also find to be particularly enriched in human TAFs in our transcriptomic analysis (Fig. S3D). Moreover, we found that TAFs are morphologically distinct from CAFs, consistent with a study of TAFs, which again referred them as “peri-tumor fibroblasts,” isolated from tongue squamous cell carcinoma and showed different cell organization in coculture compared to CAFs (45). Ba et al observed that CAFs grow in a more disorganized manner compared to peri-tumor fibroblasts. These observations are consistent with our results: we show that primary TAFs are organized in striated pattern compared to primary CAFs, which present as randomly dispersed during culture.

To date, the most promising anti-CAF therapies target the markers α-SMA and FAP but have had disappointing results, suggesting complex pro- and anti-tumor properties for CAFs (5,46). Alexeyenko et al, characterized the transcriptional landscape of normal skin and prostate fibroblasts after cell-cell contact cocultures with PC-3 tumor cells and showed that normal fibroblasts have an inhibitory effect on cancer cells. Interestingly, the authors identified *BHLHE40* and *NR4A1* as top differentially expressed genes in the cancer-inhibitory fibroblasts after coculture with cancer cells. Of note, both of these markers were exclusively upregulated in our CAFs (Fig. 1G). Our results also show a striking epithelial morphology in HCC827 cocultured with CAFs as opposed to the mesenchymal phenotype generally described (2,47,48). HCC827 formed distinct epithelial islands surrounded by CAFs and were associated with increased cell differentiation and glycosaminoglycan production. Differentiation of cancer cells is normally associated with a less proliferative and less invasive tumor behavior in the histopathological classification of solid malignancies (49). Overall, our results suggest greater anti-tumor roles for CAFs compared to TAFs.

It is unclear whether TAFs and CAFs originate from the same cell lineage. Our results highlight *LSP1* and *THSD1* as exclusively overexpressed in TAFs compared to NFs and CAFs. *THSD1* has been previously described as a marker of hematopoietic stem cell (33) and *LSP1* is a common fibrocyte marker (34). Fibrocytes are considered to be mesenchymal cells that arise from monocyte precursors with both the inflammatory features of macrophages and the tissue remodeling properties of fibroblasts (34). Our transcriptomic results also show that TAFs express significantly higher expression of MHC class I and II molecules compared to CAFs MHC class II markers are normally expressed only by professional antigen-presenting cells including macrophages (50). These results propose that TAFs may be hematopoietic-derived cells recruited at the invasive edge of the tumor as they display sine fibrocyte-like properties. However, lineage tracing analysis should be performed to confirm this hypothesis.

Previous results from our group highlighted the role of activated fibroblasts in metabolic reprogramming toward the HBP (16). Here, our work confirms that when either CAFs or TAFs are cocultured with cancer cells, they are both metabolically reprogrammed, but differently with TAFs that predominantly involve pathways associated with glycans biosynthesis. These results motivated us to expand our multi-omics approach to cover the O-glycoproteome in addition of the transcriptome and the proteome in our analysis of TAFs in the context of tumor-stroma crosstalk. Interestingly, these different methods showed a modest overlap of biological functions and unique pathways enriched in each molecular dimension. Of note, O-glycosylation implicated protein associated with the regulation of the cell cycle, cell death and metabolism. This is not surprising as several factors involved in proliferation, apoptosis and cellular energetics have been shown to be stabilized by O-glycosylation (20).

Interestingly, CDK4 was the only target differentially upregulated in TAFs, after coculture with cancer cells, across all -omics platforms used in this study (transcriptomic, proteomic, O-glycoproteomic). In depth analysis of our O-glycoproteome data revealed the importance of the CDK4-pRb axis in the stroma as both of these proteins were differentially O-glycosylated after TAFs:HCC827 coculture. CDK4 is known to phosphorylate the tumor suppressor RB, leading to its inactivation. The inactive RB releases the transcription factor E2F2, which leads to cell cycle progression and proliferation (39); notably, the transcription factor E2F2 was uniquely upregulated in TAFs, relative to the CAFs and NFs, at the gene expression level. Our study is the first to identify specific O-glycosites with their associated O-glycoforms in the CDK4-pRb axis. Wells *et al* have previously reported RB to be O-glycosylated in HeLa S3 cells at an unknown site but to our knowledge, we are the first to report CDK4 to be O-glycosylated at T153 and to identify RB O-glycosites at S534, T821 and T823. Interestingly, T153 is predicted to be located in the ATP pocket of CDK4 (37,51), which is also known as the binding site for most CDK4 inhibitors (42). Glycans can physically restrict access to a binding site or chemically modify the affinity for a target (52). The tools to validate if O-glycosylation of CDK4 at T153 affects the binding affinity of palbociclib are not readily accessible due to the limited availability of glycan antibodies and the challenges associated with generating primary fibroblast mutants. Instead, we inhibited O-glycosylation with DON and showed that the combination treatment reduces proliferation and increases senescence in both TAFs and HCC827 cocultures at a greater level than palbociclib alone. In addition, the combination treatment of DON and palbociclib promoted a CAF-like morphology in cocultured TAFs which in turn appears to reduce EMT in the HCC827. This aligns with previous studies where altering protein glycosylation mediated many cellular changes associated with EMT and cell–cell adhesion (21,53).

Moreover, the tumor suppressor role of RB has been extensively reviewed (54,55) but to our knowledge, only one group previously reported RB to be regulated by O-glycosylation. We find RB to be O-glycosylated at T821 and T823 among the three O-glycosites detected. Lentine et al showed that dephosphorylation of T821 is required for the cell to undergo apoptosis (56). In addition, these O-glycosites are located in the binding domain of the E4F1 protein, which binds to RB and indirectly stabilizes p21 (57). Although we did not observe an increase of apoptosis in the cocultured TAFs after inhibiting O-glycosylation with DON, we did observe an overall increase of p21 in the cocultured TAFs, which has been associated with senescence (43). In cancer cells, increased senescence has been reported to lead to reduced tumor proliferation but can also contribute to cancer development (58,59). As for fibroblasts, the pro- and anti-tumor roles of senescence are still being investigated as senescent fibroblasts have a different secretory profile (60). These results implicate O-glycosylation as a new avenue to regulate senescence, specifically by modulating RB interactions with E4F1 which impact p21 stabilization. However, we acknowledge that DON can also affect other metabolic enzymes using glutamine and that a site-directed mutagenesis approach to prevent O-glycosylation of CDK4 or RB1 would be preferable to confirm the role of O-glycosylation in reducing palbociclib affinity in the pRB-CDK4 signaling axis. In order to invest in developing targeted O-glycosylation pre-clinical models as opposed to traditional glycosyltransferase knockouts in mice that are generally embryonic lethal (61), we are hopeful that this O-glycoproteome dataset will help to establish the significance of specific O-glycosylation events in the context of tumor-stroma crosstalk and will set the stage for the development of future in vivo models.

In summary, we characterized the crosstalk between cancer cells and fibroblasts derived from different tumor regions including normal, adjacent and tumor core. This work placed a greater focus on an understudied subtype of fibroblasts, namely those derived from the invasive tumor front, designated here as TAFs. We found that O-glycosylation regulates the CDK4-pRB axis in TAFs cocultured with cancer cells, highlighting the relevance of the O-glycoproteome as an important dimension in modulating protein signaling through metabolic reprogramming. Lastly, modulating O-glycosylation of the CDK4-pRB axis in TAFs may indirectly decrease the mesenchymal features of cancer cells by inducing a CAF-like phenotype in TAFs. In summary, the addition of O-glycoproteome data in combination with transcriptomic and proteomic data provides new insight into the biological control of the tumor leading edge by the stroma and indicates new avenues to improve cancer treatment. We hope that these results will motivate the development of targeted preclinical models to establish the significance of specific O-glycosylation events in the lung TME and promote the importance of glycobiology in cancer.

## Acknowledgments

This work was supported by the National Cancer Institute (R25CA180993) and Les Fonds de Recherche du Québec – Santé (GB #35603 and #267646). NMR was supported by the US National Institutes of Health (K00CA212454). The authors thank Dr Parag Mallick for providing the HCC827 cell lines used in this study. We also thank Dr Sean Bendall for CyTOF reagents, Winston L Trope for providing clinical specimen information, as well as Dr Marc Driessen, Dr Yuanyuan Li, and Dr Dhanya K Nambiar for their technical support.

## Author contributions

Study Conception & Design: GB, LK, SJP and SKP; Performed Experiment and Data; Collection: GB, AB, NMR and LK; Data Analysis: GB, FJG, WZ, NMR, LK and LCM; Interpretation of data: GB, FJG, LK, LCM, SJP, AJG and SKP; Writing the first draft: GB; Fig.s Design : GB and FJG; Patient Sample Management : JAB and JBS; Supervision: JBS, CRB, SJP, AJG and SKP. All authors contributed to manuscript editing and revision.

## Data and materials availability

Further information and requests for resources and reagents should be directed to and will be fulfilled by the Lead Contact, Sylvia Plevritis. This study did not generate new unique reagents. The accession number for all data generated in this study will be established before publication and made publicly available.

## Materials and Methods

### Cell Culture

HCC827 and HCC827-eGFP NSCLC adenocarcinoma cell lines were a generous gift from Dr Parag Mallick and were grown in RPMI-1640 with L-Glutamine, supplemented with 10% fetal bovine serum (FBS), and 5% antibiotic solution (penicillin/streptomycin), at 5% CO_2_ and 37°C.

### Human Studies

Clinical aspects of this study were approved by the Stanford Institutional Review Board (IRB) in accordance with the Declaration of Helsinki guidelines for the ethical conduct of research. All patients involved provided a written informed consent. Collection and use of human tissues were approved and in compliance with data protection regulations regarding patient confidentiality (IRB protocol #15166). Following surgical resection of primary tumors from three patients at Stanford Hospital, lung adenocarcinoma specimens were immediately processed in order to establish primary fibroblast cell lines.

### Tumor and Lung Specimen Dissociation

Immediately following surgery, fresh lung adenocarcinoma matched specimens were collected from different tumor regions namely Normal Fibroblasts (NFs, >5 cm away from the tumor), Tumor-Adjacent Fibroblasts (TAFs, leading edge) and Cancer-Associated Fibroblasts (CAFs, tumor core). Samples were immersed and transferred from Stanford Hospital to the laboratory in MACS Tissue Storage Solution (Miltenyi Biotec). Then, tumors were cut into small pieces with dissecting scissors. The dissociation was performed utilizing the MACS Tumor Tissue Dissociation Kit (Miltenyi Biotec) for the tumor core samples or the Lung Dissociation Kit (Miltenyi) for the normal and adjacent tissues as per the manufacturer’s protocol. The single-cell suspensions were centrifuged at 500 × g for 5 min to pellet tumor cells, which were subsequently resuspended in RPMI-1640 and applied on a MACS SmartStrainer (70 μm, Miltenyi Biotec) for filtration Following centrifugation (500 × g, 5 minute), red blood cells were removed using the Red Blood Cell Lysis Solution according to the manufacturer’s instructions (Miltenyi Biotec). Cells were then washed, resuspended in RPMI-1640 and plated in a 35 mm dish for fibroblast expansion. At confluency, cells were trypsinized and transferred to a 25 cm^2^ flask, then split in two 75 cm^2^ flasks at confluency. Primary fibroblasts were frozen at passage 1 and all cocultures experiments were realized at passage 2.

### In Vitro Coculture Experiments and Conditioned Media Induction

HCC827 lung adenocarcinoma cells were cocultured at 1:1 ratio (8×10^5^ cells each cell type) with primary fibroblasts for 24 hours. In parallel, monocultures (16×10^6^ cells) were plated and after 24 hours, conditioned media were collected and spun down at 500 x g for 5 min. The cells were rinsed PBS 1X and treated with conditioned media from the opposite cell type for 24 hours. Then, cocultures and monoculture controls or induced with supernatant were rinsed with PBS 1X and lifted off tissue culture plates using TrypLE (Gibco) and cocultures were separated with the optimized amount of Anti-Fibroblast MicroBeads (40 μL, for up to three 100 mm Petri Dishes, Miltenyi Biotec) according to the manufacturer’s protocol to reach minimal contamination in each compartment (Fig. S6A). Cells were counted and stored accordingly at −80 °C for further analysis including flow cytometry, mass cytometry, RNA-seq, and proteomics.

Alternatively, HCC827-eGFP were cocultured at 1:1 ratio (4×10^5^ cells each) with primary TAFs for up to 72 hours. After 24 hours, cells were treated with 10 μM 6-Diazo-5-oxo-L-norleucine (DON, Sigma Aldrich), 1 μM Palbociclib (Sigma Aldrich) or a combination of both for 48 hours. Then, cocultures and monoculture controls were rinsed with PBS 1X, lifted off tissue culture plates using TrypLE (Gibco), counted, pelleted, and stored accordingly in 500 μL of flow cytometry buffer (FCB) at −80 °C for further flow cytometry analysis. Drugs were dissolved in water and non-treated controls with RPMI medium only.

### Flow Cytometry Analysis

After separation with Anti-Fibroblast MicroBeads (Miltenyi Biotec), the sorted cell compartments and monocultures were counted and aliquots of 1×10^6^ cells were prepared per condition. Aliquots were incubated for 5 min with 1 μL of Zombie Aqua™ fixable viability dye in PBS 1X, then washed with FCB (05% BSA, 002% NaN_3_, and 2 mM EDTA in PBS 1X) and centrifuged (500 × g, 5 min) before adding paraformaldehyde (PFA, EMS) at a final concentration of 16% for 10 min at room temperature in FCB. Cells were then centrifuged at 500 × g for 5 min at 4 °C to pellet cells, PFA was removed, and cells were washed again with FCB. Cells were either stored long-term at −80 °C in 500 μL of FCB or permeabilized with 100 μL of eBioscience™ Permeabilization Buffer (Invitrogen) diluted at 1X concentration for 30 min on ice with a master mix of a combination of primary antibodies (E-cadherin PE/Cy7 Biolegend, Clone 67A4, 324 115; CD10 BV421, BioLegend, Clone HI10a, 312218; Ki67 AF647, BD Biosciences, clone B56, 558615; Vimentin AF700, Novus Bio, clone RV202; α-SMA APC-Cy7, Abcore, clone SPM332; GFAT2 FITC, LSBio, clone aa175-201; FSP1 PerCP-Cy5, BioLegend, clone NJ-4F3-D1; p21 AF594, R&D Systems, clone 195720; pRB (Thr826), Invitrogen; Caspase-8 AF700, Novus Bio, clone 90A992). After the incubation with primary antibodies, cells were washed with FCB and spun down at 500 × g for 5 min at 4°C (2X). These steps were repeated with the secondary antibody as needed (IgG anti-rabbit PE, BioLegend, clone poly4064). After the last wash with FCB, cells were resuspended in 500 μL of FCB, strained and analyzed using a BD LSRFortessa™ X-20 (BD Biosciences). Results were analyzed using Cytobank single-cell analysis software with the gating strategies described in Fig. S1B, S4A and S5A depending on the analysis.

### Mass Cytometry

After separation with Anti-Fibroblast MicroBeads (Miltenyi Biotec), the sorted cell compartments and monocultures were counted and aliquots of 5×10^5^ cells were prepared per condition. To assess cell viability for mass cytometry, cell pellets were incubated for 5 min in 1 mL PBS containing cisplatin (Sigma-Aldrich) at a final concentration of 0.5 μM at room temperature. Cisplatin reaction was quenched by adding complete RPMI-1640 media (10% FBS) and subsequent centrifugation for 5 min at 500 × g. Cell pellets were resuspended in cell culture media or FCB and were fixed by adding PFA (EMS) at a final concentration of 16% for 10 min at room temperature. Cells were centrifuged at 500 × g for 5 min at 4 °C to pellet cells and remove PFA and washed once with FCB. Cell pellets were resuspended in FCB and stored at −80 °C until all conditions of the same clinical specimen were collected.

HCC827 cell lines monocultures and cocultures with NFs, TAFs, and CAFs were analyzed using mass cytometry as previously described (35). Briefly, samples were first incubated with surface antibody master mix (total volume of 100 μL) for 30 min at room temperature Samples were then washed with FCB and permeabilized with methanol for 10 min on ice. Following two washes with FCB, samples were incubated with the intracellular antibody master mix (total volume of 100 μL) for 30 min at room temperature. Samples were washed twice with FCB and resuspended in PBS containing 1:5000 191Ir/193Ir MaxPar Nucleic Acid Intercalator (Fluidigm) and 16% PFA to stain μDNA and stored at 4 °C for 1–3 days. Prior to mass cytometry analysis, cells were washed once with FCB, twice with filtered double-distilled water, and finally resuspended (~10 stained cells/mL) in filtered double-distilled water containing normalization beads (EQ Beads, Fluidigm). During event acquisition, pooled filtered cell samples were kept on ice at all times and introduced into the CyTOF 2 (Fluidigm) using the Super Sampler (Victorian Airship and Scientific Apparatus, Alamo, CA, USA). The events length, barcoding channels (102Pd, 104Pd, 105Pd, 106Pd, 108Pd, 110Pd), normalization beads (140Ce, 151Eu, 153Eu, 165Ho, 175Lu), DNA (191Ir and 193Ir), and dead cells (195Pt and 196Pt) were recorded and the full list of antibody metal isotopes recorded can be found in the key resources table.

### Mass Cytometry Data Processing

Normalization and single-cell debarcoding were performed through respective algorithms as described previously (62) and transformed using the inverse hyberbolic sine (ArcSinh) function with a cofactor of 552. Debarcoded samples were uploaded as separate FCS files and analyzed on Cytobank. Non-viable (cisplatin-positive) and apoptotic (cleaved Caspase-3) cells were removed for all subsequent single-cell analysis Cytobank software was used for traditional cytometry statistics and visualization (histograms, density plots, heatmaps) and the X-shift algorithm (63) was used in the Vortex clustering environment to visualize HCC827 cancer cells and primary fibroblasts.

### Projections onto the EMT–MET PHENOSTAMP Phenotypic Map

PHENOSTAMP was downloaded from GitHub under https://githubcom/anchangben/PHENOSTAMP. Briefly, FCS files previously gated in Cytobank according to vimentin, cytokeratins, CD104, slug, and low signaling (Fig. S7B) were uploaded into R and the PHENOSTAMP algorithm was used to project HCC827 monocultures and cocultures on the 2D EMT–MET state map as previously described (35).

### RNA Sequencing and Analysis

After in vitro coculture experiments, 1×10^5^ cells were pelleted, stored in RNA later stabilization solution (Invitrogen) and sent to MedGenome Inc (Foster City, CA, USA) for RNA extraction, library preparation, and analysis. Briefly, the RNA was extracted using a Macherey-Nagel Nucleospin analyzer (Fisher Scientific) according to the manufacturer’s protocol. Sample and library quality control were performed with Qubit (ThermoFisher) and Tapesation (Agilent) bioanalyzers. The library was prepared using Illumina TruSeq stranded mRNA and the NovaSeq PE100 sequencing run type.

Expression levels of RNA sequencing data were quantified using a quasi-mapping two phase inference algorithm implemented in Salmon (version 091) (64). Transcript sequences GRCh38 from Gencode were used for reference transcriptome indexing Transcript-per-million (TPM) value was used as normalized expression level unit. Quantified gene expressions were then log transformed for downstream analysis.

Highly variable genes with variances over 75% percentile were included for differentially expressed gene (DEG) analysis DEGs were identified between NFs, TAFs and CAFs as well as before and after coculture with HCC827 from paired patient samples DEG analysis was performed in R using paired Student’s t-test. False discovery rate was used to adjust the p values to ensure the control for multiple testing. We used two criteria for DEGs: (i) false discovery rate less than 0.05, (ii) average fold change greater than 1.5. Permutation test on two class paired samples from R package SAM (65) was also used to verify the differentially expressed genes.

### O-glycopeptide Enrichment

After in vitro coculture experiments, 1×10^6^ cells were pelleted and resuspended in 150 μL of lysis buffer (50 mM Tris, pH 8.0, 2% sodium dodecyl sulfate (SDS), 1X protease inhibitor cocktail, 25 mM NaF, 100 μM Na_3_VO_4_, 5 mM EGTA and 5 mM EDTA). Lysates were sonicated for a few seconds on ice with a Q500 sonicator at 20% intensity (Qsonica) to shear DNA/reduce viscosity, then centrifuged at 14000 rpm (4 °C, 10 min) for supernatant collection. Protein quantification was performed using a Pierce Bicinchoninic acid (BCA) protein assay kit (Thermo Scientific) and 300 μg of proteins were used for O-glycans enrichment adapted as previously described (25). Briefly, samples were reduced in 10 mM TCEP in a final volume of 50 μL for 1 h at 60°C. Samples were then alkylated with 15 mM iodoacetamide for 30-45 min at room temperature in the dark SDS was then removed by adding ice cold acetone to precipitate proteins and incubated overnight at −20 °C. Samples were vortexed and spun down at 15000 rpm at 4 °C for 15 min. Acetone was removed without disturbing the pellets and samples were left to dry for 3-5 min. The proteins were resuspended in 100 μL of 50 mM ammonium bicarbonate and sonicated until completely dissolved. Next, proteins were digested with sequencing grade modified trypsin enzyme (Thermo Fishier Scientific) in a 1:30 w:w ratio trypsin:proteins. Samples were vortexed and incubated at 37 °C for overnight without shaking. Following digestion, samples were deglycosylated with PNGase F enzyme to remove N-linked glycans and incubated overnight. The next day, samples were desalted using C18 columns (Thermo Scientific). Columns were placed on a vacuum manifold, conditioned with 3 mL of MeOH, followed by 3 mL of acetonitrile 0.1 % formic acid, and then followed by 4 mL of water with 0.1 % formic Samples were brought up to 1 mL with water and 0.1 % formic acid (Fisher Scientific) and loaded on the columns. Samples were washed with 2 mL of water with 0.1 % formic acid and eluted 2X with 750 μL of 80% acetonitrile with 0.1 % formic acid. Lastly, samples were dried using a speed vacuum (LabConco) concentrator and reconstituted in 50 μL of anhydrous DMSO for O-glycopeptide enrichment using Phenylboronic Acid (PBA) solid phase extraction First, the PBA columns were conditioned 5 x 1 mL of anhydrous DMSO and using a rubber hand pipet dropper to slowly force the liquid into the bed resin. Then, samples were loaded on the column, allowed to enter the resin bed, and then incubated for 2 hours at 37 °C. Next, samples were washed with 1 mL of acetonitrile and eluted with 750 μL of 0.1% trifluoroacetic acid in water (x2) allowing the solvent to enter the resin bed and incubated for 30 min at room temperature for each elution Samples were collected in a 2 mL vial, dried using a vacuum concentrator and resuspend in 15-20 μL of 2% acetonitrile and 0.1 % formic acid for mass spectrometry analysis.

### Mass Spectrometry Analysis

All samples were resuspended in 0.2% formic acid in water prior to LC-MS/MS analysis. Protein samples were resuspended at 1 μg/μL with 1 μg injected on column, while enriched glycopeptides were resuspended in 20 μL total with 4 μL injected per analysis. Tryptic (glyco)peptide mixtures were separated over a 25 cm EasySpray reversed phase LC column (75 μm inner diameter packed with 2 μm, 100 Å, PepMap C18 particles, Thermo Fisher Scientific). The mobile phases (A: water with 0.2% formic acid and B: acetonitrile with 0.2% formic acid) were driven and controlled by a Dionex Ultimate 3000 RPLC nano system (Thermo Fisher Scientific). An integrated loading pump was used to load peptides onto a trap column (Acclaim PepMap 100 C18, 5 μm particles, 20 mm length, Thermo Fisher Scientific) at 8 μL/minute, which was put in line with the analytical column 4 minutes into the gradient for the total protein samples. The gradient increased from 0% to 5% B over the first 4 min of the analysis, followed by an increase from 5% to 25% B from 6 to 72 min, an increase from 25% to 90% B from 72 to 74 min, isocratic flow at 90% B from 74 to 78 min, and a re-equilibration at 0% for 12 min for a total analysis time of 90 min. For the glycopeptide mixtures, the trap column was loaded at 5 μL/min and put in line with analytical column at 6 min. The gradient increased from 0% to 5% B over the first 6 min of the analysis, followed by an increase from 5% to 25% B from 6 to 68 min, an increase from 25% to 45% B from 68 to 72 min, an increase from 45% to 90% B from 72 to 74 min, isocratic flow at 90% B from 74 to 78 min, and a re-equilibration at 0% for 12 min for a total analysis time of 90 min.

Eluted peptides were analyzed on an Orbitrap Fusion Tribrid MS system (Thermo Fisher Scientific). Precursors were ionized using an EASY-Spray ionization source (Thermo Fisher Scientific) source held at +22 kV compared to ground, and the column was held at 40 °C. The inlet capillary temperature was held at 275 °C. For the total protein peptide mixtures, survey scans of peptide precursors were collected in the Orbitrap from 350-1350 m/z with an AGC target of 1,000,000, a maximum injection time of 50 ms, and a resolution of 60,000 at 200 m/z. Monoisotopic precursor selection was enabled for peptide isotopic distributions, precursors of z = 2-5 were selected for data-dependent MS/MS scans for 1 second of cycle time, and dynamic exclusion was set to 30 s with a ±10 ppm window set around the precursor monoisotope. An isolation window of 1 m/z was used to select precursor ions with the quadrupole MS/MS scans were collected using HCD at 30 normalized collision energy with an AGC target of 30,000 and a maximum injection time of 22 ms. Mass analysis was performed in the linear ion trap using the “Turbo” scan speed while scanning from 120-1420 Th. Glycopeptide analyses were performed using product-dependent triggering as previously described (66). Here, survey scans of peptide precursors were collected in the Orbitrap from 500-1800 m/z with an AGC target of 400,000, a maximum injection time of 50 ms, RF lens at 60%, and a resolution of 120,000 at 200 m/z. Monoisotopic precursor selection was enabled for peptide isotopic distributions, precursors of z = 2-8 were selected for data-dependent MS/MS scans for 3 s of cycle time, and dynamic exclusion was set to 60 s with a ±10 ppm window set around the precursor monoisotope. Precursor priorities were set to favor highest charge state and lowest m/z precursor ions, and an isolation window of 2 m/z was used to select precursor ions with the quadrupole. In the product-dependent scheme, the presence of nine oxonium ions (126055, 1380549, 1440655, 1680654, 186076, 2040865, 2740921, 2921027, and 3661395) in “scouting” higher-energy collisional dissociation (HCD) MS/MS scan triggered acquisition of a second MS/MS scan (67–69). The “scout HCD” scan used an automated scan range determination and a first mass of 100 Th, a normalized collision energy (nce) of 25, an AGG target value of 50,000, a maximum injection time of 60 ms, and a resolution of 30,000 at 200 m/z. If at least two of the nine listed oxonium ions were present in the scout HCD scan within a ±10 ppm tolerance and were among the 20 most intense peaks, a second MS/MS scan was triggered. This second MS/MS scan had a fixed scan range of 120-4,000 m/z (this extended mass range has been shown to improve glycopeptide characterization (70)), an AGC target of 100,000 ions, a maximum injection time of 200 ms, and a resolution of 30,000 at 200 m/z. Calibrated charge dependent parameters for calculating reagent AGC targets and ion-ion reaction times (71) were enabled for EThcD scans with a supplemental activation 25 nce.

### Peptide identification, quantification, data processing and statistical analysis

The resulting MS raw data files were searched using Byonic software (Protein Metrics) (72). For total protein samples, data were searched using the entire human proteome downloaded from Uniprot (73) (reviewed, 20428 entries), with decoys appended within the Byonic environment as reversed sequences. Cleavage specificity was set as fully specific for C-terminal to R and K residues (fully tryptic) with two missed cleavages allowed, and fragmentation type was set to QTOF/HCD. Precursor mass tolerance was set to 10 ppm with fragment mass tolerance set to 0.4 Da The total common max value for modifications was set to 3 and the total rare max was set to 1. Modifications used were carbamidomethyl at cysteine (+57,021644, fixed), oxidation at methionine (+15994915, common2), and acetylation at the protein N-temrinal (+42010565, rare1). The protein FDR cutoff was set to 1% Unique databases of identified proteins were created for each total protein sample file using the “Create focused database” feature in Byonic. Following searching of all total protein runs, identified proteins in all focused database files were combined into one fasta file with 4,495 non-redundant entries to make an appropriately sized database for glycopeptide searching (74). Using this database, glycopeptide raw files were searched in Byonic using a glycan database of 9 common O-glycans provided by Byonic, with the glycan modifications being denoted as common2 in Byonic (meaning they could each occur twice in a sequence). The total common max value was set to 3 and the total rare max was set to 1. Other modifications were: carbamidomethyl at cysteine (+57,021644, fixed), oxidation at methionine (+15994915, common2), acetylation at the protein N-terminal (+42010565, rare1), and deamidated asparagine (+0984016, rare1). Cleavage specificity was set as fully specific for C-terminal to R and K residues (fully tryptic) with three missed cleavages allowed. Precursor mass tolerance was set to 10 ppm with fragment mass tolerance(s) set to 20 ppm for both HCD and EThcD, and protein FDR was set to 1% was used that was built using the “Create focused database” feature in Byonic for standard shotgun proteomic experiments on unenriched tryptic peptides from HEK293 whole cell lysate.

Quantitative information was extracted from MS1 spectra of all identified peptides using an in-house R script based on MSnbase package (30) as the AUC of the extracted ion current (XIC) of all remaining peptides after the alignment of the chromatographic runs. Then protein abundances changes were analyzed using the Generic Integration Algorithm. All quantitative information is expressed in terms of Z-scores at protein or at peptide level, according to (75) in which the log2 ratios, were calculated comparing the AUC of peptides in TAFs cocultures against the average of the controls; are pondered using the corresponding statistical weight, calculated at spectrum level, according to WSPP model; rescaled, and standardized to a normal distribution N(0,1). The validity of the null hypothesis at each level (spectrum, peptide, and protein) was carefully checked by plotting the cumulative distributions. The variances at the scan, peptide and protein levels, as well as protein expression changes were determined only with non-modified peptides. Afterwards, in the intact glycoproteomics analysis the glycan-containing peptides were included in the analysis to determine the ones deviating more than expected from the rest of peptides (non-modified) belonging to the same protein, applying the same method used in (75). The final statistical comparison was performed using Student’s-t test.

### Immunofluorescence

TAFs:HCC827-eGFP cocultures and HCC827-eGFP monocultures were grown on coverslip using the same culture conditions previously described. Coverslips were rinsed 3 times with a cold solution of 5% BSA diluted in PBS and cells were fixed for 10 min in PFA 4% diluted in PBS. Then, the coverslips were incubated in 0.1% Triton X-100 for 10 min, rinsed with PBS, blocked with 10% goat serum diluted in PBS for 30 min and incubated with the CD44 antibody (Cell Signaling), cat#3570) in blocking solution in a humidified chamber overnight at 4 °C. After three washes in PBS, the coverslips were incubated with the Cy3-conjugated goat anti-rabbit IgG (Abcam, ab97035) in blocking solution for 1 hour at RT in dark. The coverslips were intensively washed with PBS and mounted on slides with a drop of mounting medium containing DAPI (Invitrogen). The sections were observed under the BZ-X800 fluorescence microscope (Keyence, IL, USA).

### Quantification and statistical analysis

Statistical testing for PHENOSTAMP results were performed with GraphPad Prism Software 8.4, using two-way ANOVA followed by multiple comparisons Fisher least significant difference (LSD) test as indicated in figure legends. Error bars in the figures represent the SD Statistical significance: *p <0.05; **p <0.01. Gene set enrichment analysis overlap and activity z-scores p values have been calculated using the Ingenuity Pathway Analysis Software (Qiagen) using the previously described method (76). Metabolic pathway gene overlaps were calculated using a hypergeometric test. Statistical analysis of RNA-seq and mass spectrometry data is described in their respective method sections.

## References

1. Deryugina EI, Kiosses WB, Jolla L, Facility MC, Jolla L. Intratumoral Cancer Cell Intravasation can occur Independent of Invasion into the Adjacent Stroma. Cell Rep. 2017;19:601–16.

2. Kalluri R. The biology and function of fibroblasts in cancer. Nat Publ Gr. 2016;16.

3. Ishii G, Ochiai A, Neri S. Phenotypic and functional heterogeneity of cancer-associated fibroblast within the tumor microenvironment. Adv Drug Deliv Rev [Internet]. Elsevier B.V.; 2016;99:186–96. Available from: http://dx.doi.org/10.1016/j.addr.2015.07.007

4. Bu L, Baba H, Yoshida N, Miyake K, Yasuda T, Uchihara T, et al. Biological heterogeneity and versatility of cancer-associated fibroblasts in the tumor microenvironment. Oncogene [Internet]. Springer US; 2019; Available from: http://www.nature.com/articles/s41388-019-0765-y

5. Özdemir BC, Pentcheva-Hoang T, Carstens JL, Zheng X, Wu CC, Simpson TR, et al. Depletion of carcinoma-associated fibroblasts and fibrosis induces immunosuppression and accelerates pancreas cancer with reduced survival. Cancer Cell. 2014;25:719–34.

6. Lambrechts D, Wauters E, Boeckx B, Aibar S, Nittner D, Burton O, et al. Phenotype Moulding of Stromal Cells in the Lung Tumour Microenvironment. Nature. 2018;1–23.

7. Elyada E, Bolisetty M, Laise P, Flynn WF, Courtois ET, Burkhart RA, et al. Cross-species single-cell analysis of pancreatic ductal adenocarcinoma reveals antigen-presenting cancer-associated fibroblasts. Cancer Discov. 2019;9:1102–23.

8. Zhao Z, Xiong S, Wang R, Li Y, Wang X, Wang Y, et al. Peri-tumor fibroblasts promote tumorigenesis and metastasis of hepatocellular carcinoma via Interleukin6/STAT3 signaling pathway. Cancer Manag Res. 2019;11:2889–901.

9. Sivridis E, Giatromanolaki A, Koukourakis MI. Proliferating fibroblasts at the invading tumour edge of colorectal adenocarcinomas are associated with endogenous markers of hypoxia, acidity, and oxidative stress. J Clin Pathol. 2005;58:1033–8.

10. Costa A, Kieffer Y, Scholer-Dahirel A, Pelon F, Bourachot B, Cardon M, et al. Fibroblast Heterogeneity and Immunosuppressive Environment in Human Breast Cancer. Cancer Cell [Internet]. Elsevier; 2018;33:463–479.e10. Available from: http://dx.doi.org/10.1016/j.ccell.2018.01.011

11. Manousopoulou A, Hayden A, Mellone M, Baquero DJG, White CH, Noble F, et al. Quantitative proteomic profiling of primary cancer-associated fibroblasts in oesophageal adenocarcinoma: CAF proteomic profiling in OAC. Br J Cancer [Internet]. 2018;118:1200–1207. Available from: https://eprints.soton.ac.uk/417703/

12. Ligorio M, Sil S, Malagon-Lopez J, Nieman LT, Misale S, Di Pilato M, et al. Stromal Microenvironment Shapes the Intratumoral Architecture of Pancreatic Cancer. SSRN Electron J [Internet]. Elsevier; 2018;178:160–175.e27. Available from: http://dx.doi.org/10.1016/j.cell.2019.05.012

13. Gillette MA, Satpathy S, Cao S, Dhanasekaran SM, Vasaikar S V., Krug K, et al. Proteogenomic Characterization Reveals Therapeutic Vulnerabilities in Lung Adenocarcinoma. Cell. 2020;182:200–225.e35.

14. Xu JY, Zhang C, Wang X, Zhai L, Ma Y, Mao Y, et al. Integrative Proteomic Characterization of Human Lung Adenocarcinoma. Cell [Internet]. Elsevier; 2020;182:245–261.e17. Available from: http://dx.doi.org/10.1016/j.cell.2020.05.043

15. Li C, Sun Y Di, Yu GY, Cui JR, Lou Z, Zhang H, et al. Integrated Omics of Metastatic Colorectal Cancer. Cancer Cell [Internet]. Elsevier Inc.; 2020;38:734–747.e9. Available from: https://doi.org/10.1016/j.ccell.2020.08.002

16. Zhang W, Bouchard G, Yu A, Shafiq M, Jamali M, Shrager JBJB, et al. GFPT2-expressing cancer-associated fibroblasts mediate metabolic reprogramming in human lung adenocarcinoma. Cancer Res. 2018;78:3445–57.

17. Kim J, Lee HM, Cai F, Ko B, Yang C, Lieu EL, et al. The hexosamine biosynthesis pathway is a targetable liability in KRAS/LKB1 mutant lung cancer. Nat Metab [Internet]. Springer US; 2020;2. Available from: http://dx.doi.org/10.1038/s42255-020-00316-0

18. National Research Council. Transforming glycoscience: A roadmap for the future. 2012.

19. Steentoft C, Vakhrushev SY, Joshi HJ, Kong Y, Vester-Christensen MB, Schjoldager KT-BG, et al. Precision mapping of the human O-GalNAc glycoproteome through SimpleCell technology. EMBO J [Internet]. 2013;32:1478–88. Available from: http://www.pubmedcentral.nih.gov/articlerender.fcgi?artid=3655468&tool=pmcentrez&rendertype=abstract

20. Peixoto A, Relvas-Santos M, Azevedo R, Lara Santos L, Ferreira JA. Protein glycosylation and tumor microenvironment alterations driving cancer hallmarks. Front Oncol. 2019;14:380.

21. Phillips RM, Lam C, Wang H, Tran PT. Bittersweet tumor development and progression: Emerging roles of epithelial plasticity glycosylations [Internet]. 1st ed. Adv. Cancer Res. Elsevier Inc.; 2019. Available from: http://dx.doi.org/10.1016/bs.acr.2019.01.002

22. Cummings RD. The repertoire of glycan determinants in the human glycome. Mol Biosyst. 2009;5:1087–104.

23. Huang J-M, Wu M-X, Zhang Y, Kong S-Y, Liu M-Q, Jiang B-Y, et al. OGP: A Repository of Experimentally Characterized O-Glycoproteins to Facilitate Studies on O-Glycosylation. bioRixv. 2020;

24. Woo CM, Iavarone AT, Spiciarich DR, Palaniappan KK, Bertozzi CR. Isotope-targeted glycoproteomics (IsoTaG): a mass-independent platform for intact N- and O-glycopeptide discovery and analysis. Nat Methods [Internet]. 2015;12:561–7. Available from: http://www.nature.com/doifinder/10.1038/nmeth.3366

25. Wang X, Yuan Z-F, Fan J, Karch KR, Ball LE, Denu JM, et al. A Novel Quantitative Mass Spectrometry Platform for Determining Protein O-GlcNAcylation Dynamics. Mol Cell Proteomics [Internet]. 2016;15:2462–75. Available from: http://www.mcponline.org/content/15/7/2462.abstract

26. Shon DJ, Malaker SA, Pedram K, Yang E, Krishnan V, Dorigo O, et al. An enzymatic toolkit for selective proteolysis, detection, and visualization of mucin-domain glycoproteins. Proc Natl Acad Sci U S A. 2020;117:21299–307.

27. Yang W, Ao M, Hu Y, Li QK, Zhang H. Mapping the O◻glycoproteome using site◻specific extraction of O◻linked glycopeptides (EXoO). Mol Syst Biol. 2018;14:1–12.

28. Malaker SA, Pedram K, Ferracane MJ, Bensing BA, Krishnan V, Pett C, et al. The mucin-selective protease StcE enables molecular and functional analysis of human cancer-associated mucins. Proc Natl Acad Sci U S A. 2019;116:7278–87.

29. Steentoft C, Vakhrushev SY, Vester-Christensen MB, Schjoldager KTBG, Kong Y, Bennett EP, et al. Mining the O-glycoproteome using zinc-finger nuclease-glycoengineered SimpleCell lines. Nat Methods. 2011;8:977–82.

30. Gatto L, Lilley KS. Msnbase-an R/Bioconductor package for isobaric tagged mass spectrometry data visualization, processing and quantitation. Bioinformatics. 2012;28:288–9.

31. Gao Z, Shi R, Yuan K, Wang Y. Expression and prognostic value of E2F activators in NSCLC and subtypes: a research based on bioinformatics analysis. Tumor Biol [Internet]. Tumor Biology; 2016;37:14979–87. Available from: http://dx.doi.org/10.1007/s13277-016-5389-z

32. Kent LN, Leone G. The broken cycle: E2F dysfunction in cancer. Nat Rev Cancer [Internet]. Springer US; 2019;19:326–38. Available from: http://dx.doi.org/10.1038/s41568-019-0143-7

33. Takayanagi SI, Hiroyama T, Yamazaki S, Nakajima T, Morita Y, Usui J, et al. Gene Expression Profiles of Non-Small Cell Lung Cancer: Survival Prediction and New Biomarkers. Blood. 2006;107:4317–25.

34. Reilkoff RA, Bucala R, Herzog EL. Fibrocytes: emerging effector cells in chronic inflammation. Nat Rev Immunol. 2011;11:427–435.

35. Karacosta LG, Anchang B, Ignatiadis N, Kimmey SC, Benson JA, Shrager JB, et al. Mapping Lung Cancer Epithelial-Mesenchymal Transition States and Trajectories with Single-Cell Resolution. Nat Commun [Internet]. 2019;10:5587. Available from: http://biorxiv.org/content/early/2019/03/07/570341.abstract

36. Brockhausen I, Stanley P. Chapter 10 O-GalNAc Glycans. Essentials Glycobiol [Internet]. 2017;1–9. Available from: https://www.ncbi.nlm.nih.gov/books/NBK453030/

37. Echalier A, Hole AJ, Lolli G, Endicott JA, Noble MEM. An inhibitor’s-eye view of the atp-binding site of CDKs in different regulatory states. ACS Chem Biol. 2014;9:1251–6.

38. Pandey K, An HJ, Kim SK, Lee SA, Kim S, Lim SM, et al. Molecular mechanisms of resistance to CDK4/6 inhibitors in breast cancer: A review. Int J Cancer. 2019;145:1179–88.

39. Klein ME, Kovatcheva M, Davis LE, Tap WD, Koff A. CDK4/6 Inhibitors: The Mechanism of Action May Not Be as Simple as Once Thought. Cancer Cell [Internet]. Elsevier; 2018;34:9–20. Available from: http://dx.doi.org/10.1016/j.ccell.2018.03.023

40. Chen W, Do KC, Saxton B, Leng S, Filipczak P, Tessema M, et al. Inhibition of the hexosamine biosynthesis pathway potentiates cisplatin cytotoxicity by decreasing BiP expression in non-small-cell lung cancer cells. Mol Carcinog [Internet]. 2019; Available from: http://doi.wiley.com/10.1002/mc.22992

41. Naz S, Sowers A, Choudhuri R, Wissler M, Gamson J, Mathias A, et al. Abemaciclib, a selective CDK4/6 inhibitor, enhances the radiosensitivity of non–small cell lung cancer in vitro and in vivo. Clin Cancer Res. 2018;24:3994–4005.

42. Goel S, DeCristo MJ, McAllister SS, Zhao JJ. CDK4/6 Inhibition in Cancer: Beyond Cell Cycle Arrest. Trends Cell Biol [Internet]. Elsevier Ltd; 2018;28:911–25. Available from: https://doi.org/10.1016/j.tcb.2018.07.002

43. Ashraf HM, Moser J, Spencer SL. Senescence Evasion in Chemotherapy: A Sweet Spot for p21. Cell [Internet]. Elsevier Inc.; 2019;178:267–9. Available from: https://doi.org/10.1016/j.cell.2019.06.025

44. Qin X, Sufi J, Vlckova P, Kyriakidou P, Acton SE, Li VSW, et al. Cell-type-specific signaling networks in heterocellular organoids. Nat Methods [Internet]. Springer US; 2020;17:335–42. Available from: http://dx.doi.org/10.1038/s41592-020-0737-8

45. Ba P, Zhang X, Yu M, Li L, Duan X, Wang M, et al. Cancer associated fibroblasts are distinguishable from peri-tumor fibroblasts by biological characteristics in TSCC. Oncol Lett. 2019;18:2484–90.

46. Nurmik M, Ullmann P, Rodriguez F, Haan S, Letellier E. In search of definitions: Cancer-associated fibroblasts and their markers. Int J Cancer. 2020;146:895–905.

47. Yu Y, Xiao C-H, Tan L-D, Wang Q-S, Li X-Q, Feng Y-M. Cancer-associated fibroblasts induce epithelial–mesenchymal transition of breast cancer cells through paracrine TGF-β signalling. Br J Cancer. 2014;110:724–32.

48. Luo M, Luo Y, Mao N, Huang G, Teng C, Wang H, et al. Cancer-Associated Fibroblasts Accelerate Malignant Progression of Non-Small Cell Lung Cancer via Connexin 43-Formed Unidirectional Gap Junctional Intercellular Communication. Cell Physiol Biochem [Internet]. 2018;51:315–36. Available from: http://www.ncbi.nlm.nih.gov/pubmed/30453281

49. Weng CF, Huang CJ, Huang SH, Wu MH, Tseng AH, Sung YC, et al. New international association for the study of lung cancer (Iaslc) pathology committee grading system for the prognostic outcome of advanced lung adenocarcinoma. Cancers (Basel). 2020;12:1–14.

50. Roche PA, Furuta K. The ins and outs of MHC class II-mediated antigen processing and presentation. Nat Rev Immunol [Internet]. Nature Publishing Group; 2015;15:203–16. Available from: http://dx.doi.org/10.1038/nri3818

51. Shafiq MI, Steinbrecher T, Schmid R. Fascaplysin as a specific inhibitor for cdk4: Insights from molecular modelling. PLoS One. 2012;7:1–9.

52. Pinho SS, Reis CA. Glycosylation in cancer: Mechanisms and clinical implications. Nat Rev Cancer [Internet]. Nature Publishing Group; 2015;15:540–55. Available from: http://dx.doi.org/10.1038/nrc3982

53. Taparra K, Tran PT, Zachara NE. Hijacking the Hexosamine Biosynthetic Pathway to Promote EMT-Mediated Neoplastic Phenotypes. Front Oncol. Frontiers Media SA; 2016;6:85.

54. Burkhart DL, Sage J. Cellular mechanisms of tumour suppression by the retinoblastoma gene. Nat Rev Cancer. 2008;8:671–82.

55. Giacinti C, Giordano A. RB and cell cycle progression. Oncogene. 2006;25:5220–7.

56. Lentine B, Antonucci L, Hunce R, Edwards J, Marallano V, Krucher NA. Dephosphorylation of threonine-821 of the retinoblastoma tumor suppressor protein (Rb) is required for apoptosis induced by UV and Cdk inhibition. Cell Cycle. 2012;11:3324–30.

57. Fajas L, Paul C, Zugasti O, Le Cam L, Polanowska J, Fabbrizio E, et al. pRB binds to and modulates the transrepressing activity of the E1A-regulated transcription factor p120E4F. Proc Natl Acad Sci U S A. 2000;97:7738–43.

58. Gewirtz D a. Autophagy and senescence in cancer therapy. J Cell Physiol [Internet]. 2014 [cited 2014 Dec 25];229:6–9. Available from: http://www.ncbi.nlm.nih.gov/pubmed/23794221

59. Wang B, Kohli J, Demaria M. Senescent Cells in Cancer Therapy: Friends or Foes? Trends in Cancer [Internet]. The Author(s); 2020;6:838–57. Available from: https://doi.org/10.1016/j.trecan.2020.05.004

60. Coppé JP, Desprez PY, Krtolica A, Campisi J. The senescence-associated secretory phenotype: The dark side of tumor suppression. Annu Rev Pathol Mech Dis. 2010;5:99–118.

61. Widomska Justyna. What have we learned from glycosyltransferase knockouts in mice? Physiol Behav. 2017;176:139–48.

62. Zunder ER, Finck R, Behbehani GK, Amir EAD, Krishnaswamy S, Gonzalez VD, et al. Palladium-based mass tag cell barcoding with a doublet-filtering scheme and single-cell deconvolution algorithm. Nat Protoc. Nature Publishing Group; 2015;10:316–33.

63. Samusik N, Good Z, Spitzer MH, Davis KL, Nolan GP. Automated mapping of phenotype space with single-cell data. Nat Methods. 2016;13:493–6.

64. Patro R, Duggal G, Love MI, Irizarry RA, Kingsford C. Salmon provides fast and bias-aware quantification of transcript expression. Nat Methods. Nature Publishing Group; 2017;14:417–9.

65. Tusher VG, Tibshirani R, Chu G. Significance analysis of microarrays applied to the ionizing radiation response. Proc Natl Acad Sci U S A. 2001;98:5116–21.

66. Riley NM, Malaker SA, Driessen MD, Bertozzi CR. Optimal Dissociation Methods Differ for N- A nd O-Glycopeptides. J Proteome Res. 2020;19:3286–301.

67. Saba J, Dutta S, Hemenway E, Viner R. Increasing the Productivity of Glycopeptides Analysis by Using Higher-Energy Collision Dissociation-Accurate Mass-Product-Dependent Electron Transfer Dissociation. Int J Proteomics. 2012;2012:1–7.

68. Wu SW, Pu TH, Viner R, Khoo KH. Novel LC-MS2 product dependent parallel data acquisition function and data analysis workflow for sequencing and identification of intact glycopeptides. Anal Chem. 2014;86:5478–86.

69. Singh C, Zampronio CG, Creese AJ, Cooper HJ. Higher energy collision dissociation (HCD) product ion-triggered electron transfer dissociation (ETD) mass spectrometry for the analysis of N-linked glycoproteins. J Proteome Res. 2012;11:4517–25.

70. Čaval T, Zhu J, Heck AJR. Simply Extending the Mass Range in Electron Transfer Higher Energy Collisional Dissociation Increases Confidence in N-Glycopeptide Identification. Anal Chem. 2019;91:10401–6.

71. Rose CM, Rush MJP, Riley NM, Merrill AE, Kwiecien NW, Holden DD, et al. A Calibration Routine for Efficient ETD in Large-Scale Proteomics. J Am Soc Mass Spectrom. 2015;26:1848–57.

72. Bern M, Kil YJ, Becker C. Byonic: Advanced peptide and protein identification software. Curr Protoc Bioinforma. 2012;1–17.

73. Bateman A. UniProt: A worldwide hub of protein knowledge. Nucleic Acids Res. Oxford University Press; 2019;47:D506–15.

74. Khatri K, Klein JA, Zaia J. Use of an informed search space maximizes confidence of site-specific assignment of glycoprotein glycosylation. Anal Bioanal Chem. 2017;409:607–18.

75. Navarro P, Trevisan-Herraz M, Bonzon-Kulichenko E, Núñez E, Martínez-Acedo P, Pérez-Hernández D, et al. General statistical framework for quantitative proteomics by stable isotope labeling. J Proteome Res. 2014;13:1234–47.

76. Krämer A, Green J, Pollard J, Tugendreich S. Causal analysis approaches in ingenuity pathway analysis. Bioinformatics. 2014;30:523–30.

