## Supplementary Figures and Legends for "Multi-Omics Analysis of Fibroblasts from the Invasive Tumor Edge Reveals that Tumor-Stroma Crosstalk Induces O-glycosylation of the CDK4-pRB Axis"

**This PDF file includes:**

Figures S1 to S7

**Other Supplementary Materials for this manuscript include the following:**

Key Resource Table S1

Supplementary Data S1 to S6

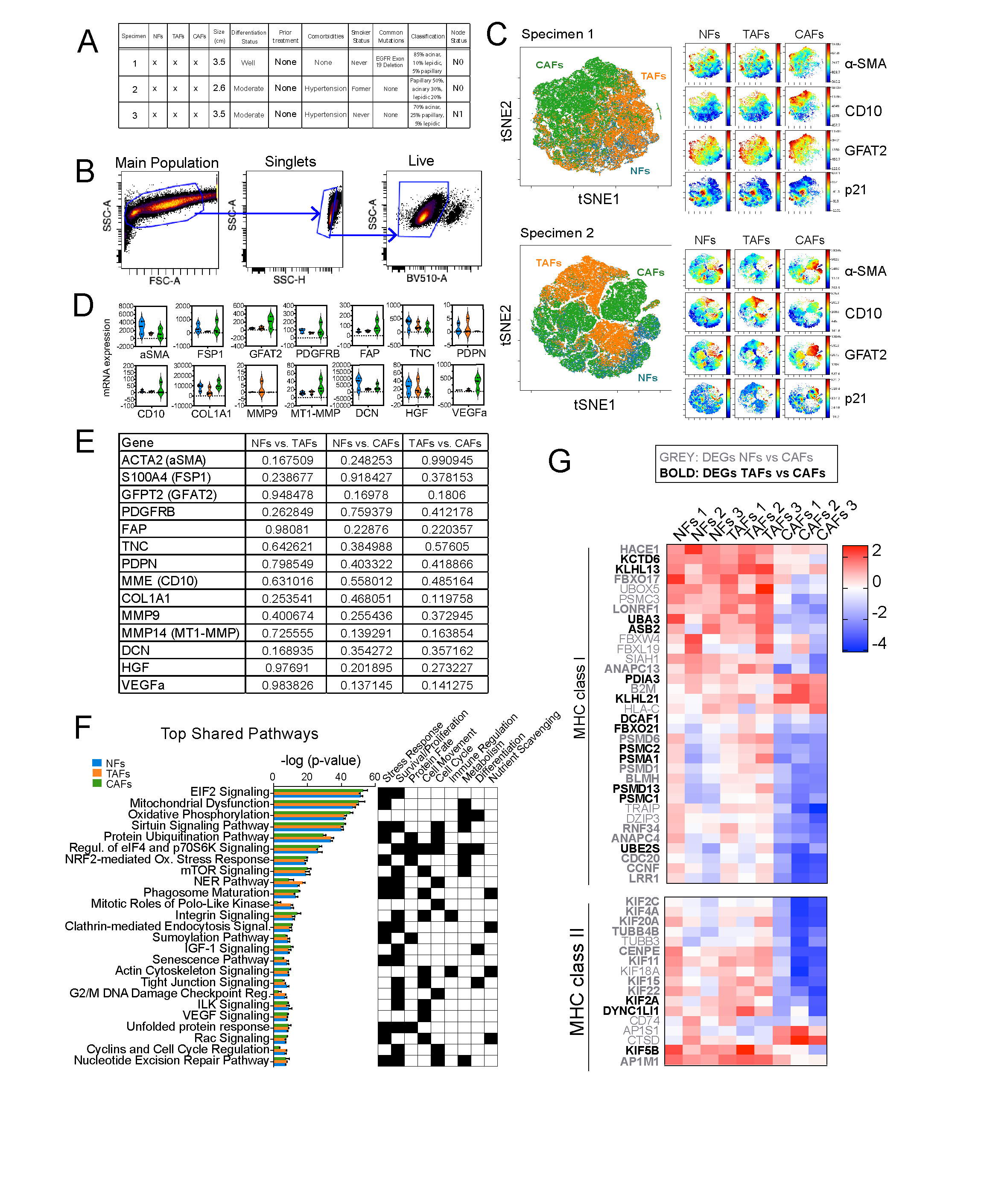

**Figure S1. Properties of LUAD fibroblast subtypes. A,** Clinical annotations and histopathological information of the three LUAD specimens analyzed in this study. **B,** Gating strategy for primary fibroblast cell lines removing debris, doublets and non-viable cells. **C,** Biological replicates representing viSNE analysis of NFs, TAFs, and CAFs with α-SMA, FSP1, GFAT2, CD10, p21, and vimentin as clustering markers and analyzed with Cytobank software. **D,** Violin plots representing mRNA expression of CAFs canonical markers measured in NFs, TAFs and CAFs and **(E)** associated p values for each group comparison. **F,** Bar graph showing the top shared pathways between NFs, TAFs and CAFs analyzed with the IPA software (Qiagen). **G,** Heatmap showing relative normalized gene expression of DEGs associated with MHC class I and II in NFs, TAFs and CAFs.

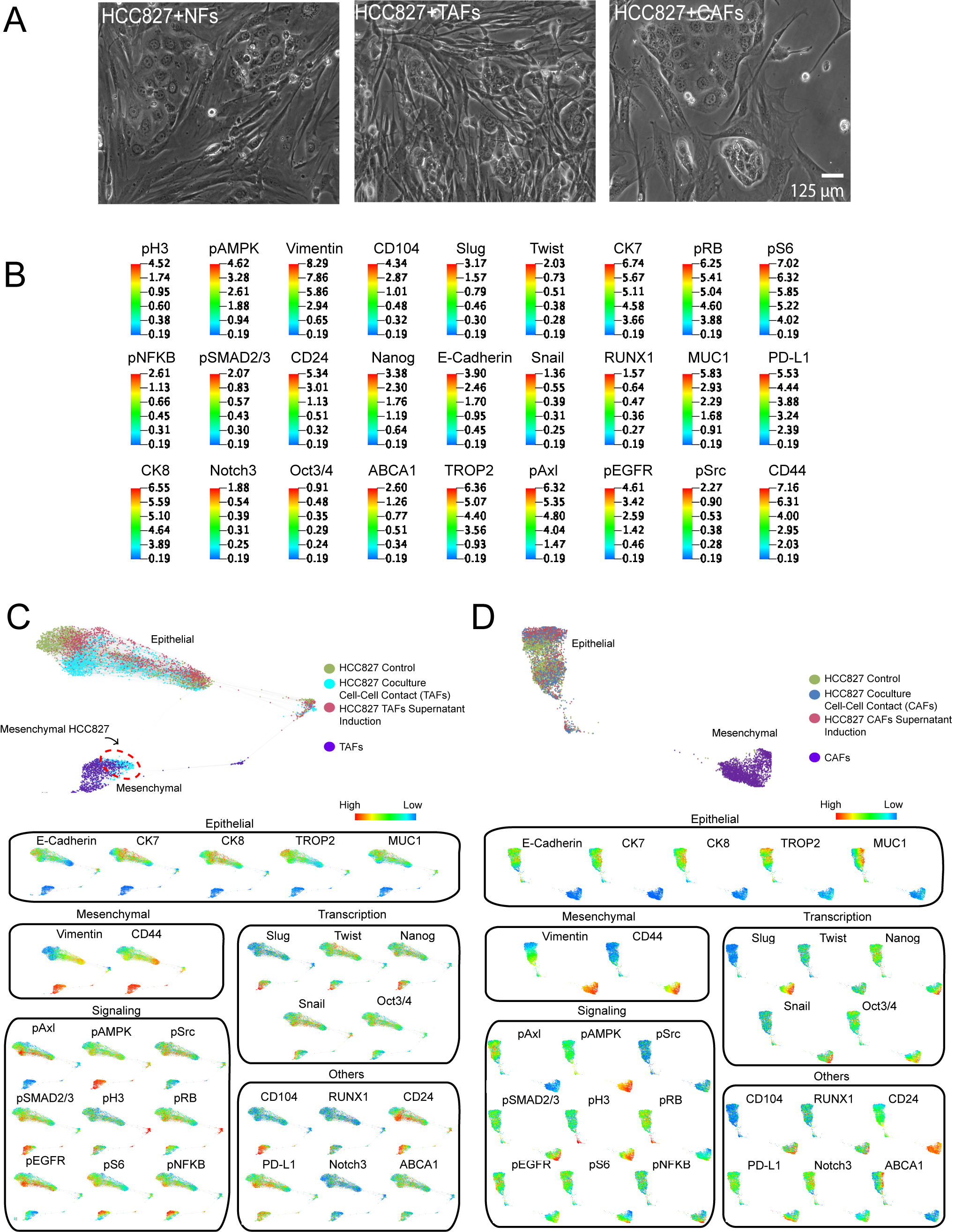

**Figure S2. Cocultures of LUAD primary fibroblasts and HCC827 analyzed with CyTOF. A,** Phase contrast microscopy images of NFs, TAFs and CAFs cocultures with HCC827 lung adenocarcinoma cell line. **B,** Scale of indicated markers measured in the HCC827 single-cell force-directed layout (arcsinh transformed data of specimen #3 presented at Figure 3C. **C,** Single-cell force-directed layout colored by protein expression of indicated markers of HCC827 cocultures with TAFs using X-shift clustering (arcsinh transformed data of specimen #2). **D,** Single-cell force-directed layout colored by protein expression of indicated markers of HCC827 cocultures with CAFs using X-shift clustering (arcsinh transformed data of specimen #1).

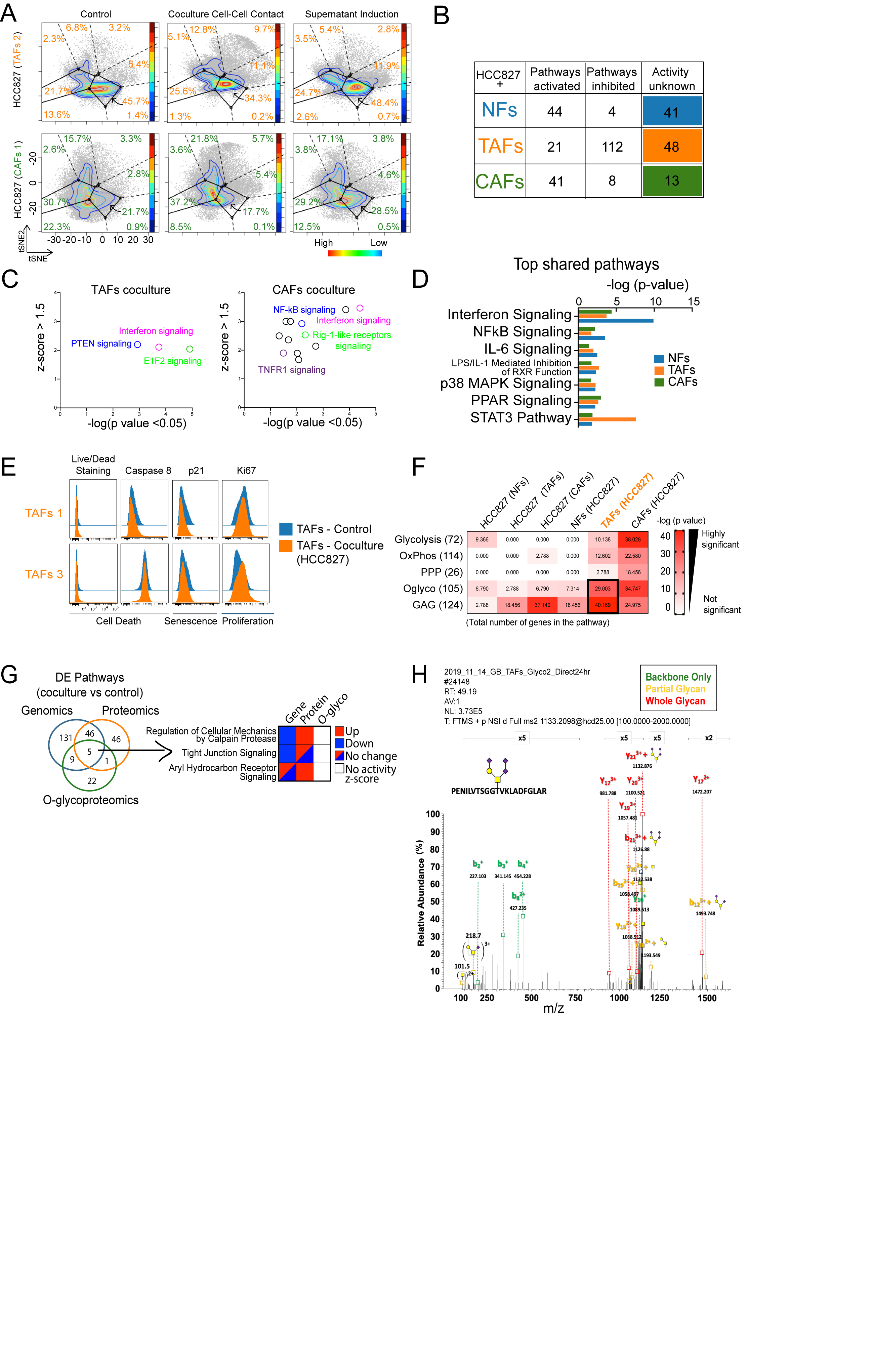

**Figure S3. Cell-cell contact cocultures promote EMT in HCC827 and metabolic reprogramming in TAFs. A,** Projections of the HCC827 monoculture controls and matched cocultures after cell-cell contact or supernatant induction with NFs, TAFs and CAFs onto the EMT–MET PHENOSTAMP phenotypic map. **B,** Pathways activated, inhibited or with unknown activity (no activity z-score available), **(C)** plots of the most significantly activated pathways in TAFs and CAFs and **(D)** top shared pathways between NFs, TAFs and CAFs after coculture with HCC827 calculated with the IPA software (Qiagen). **E,** Histograms of TAFs mono- and cocultures showing no significant difference in cell death as measured with live/dead fixable staining and caspase 8 marker using flow cytometry analysis. **F,** Heatmap showcasing the number of overlap genes between DEGs in each condition within each metabolic pathway. Color of heatmap represents -log(p-value) of the gene overlap calculated using a hypergeometric test. The number in parentheses show the total number of genes from each MSigDB gene set and tile numbers show the number of -log(p value) for each condition. **G,** Venn Diagram showing the overlap of the DE pathways identified in TAFs cocultures (relative to control monoculture) and highlighting examples of inconsistent changes across different methods of analysis. **H,** Representative HCD spectrum that provide confident localization of one glycosite in the CDK4 peptide sequence PENILVTSGGTVKLADFGLAR.

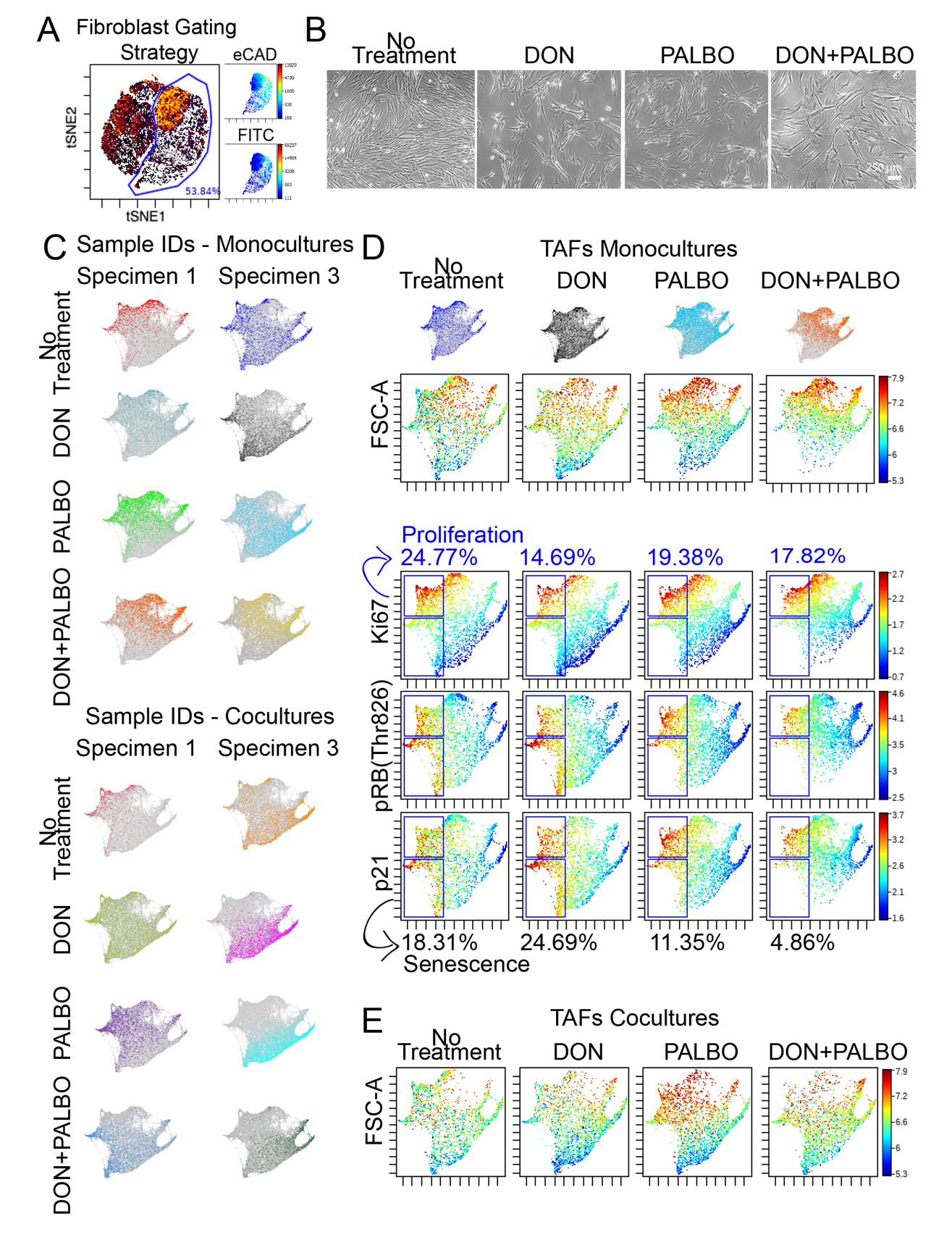

**Figure S4. Treatment with DON modulates Palbociclib efficacy in TAFs.**

**A,** Gating strategy for TAFs. After removing debris, doublets and non-viable cells as shown in Fig. S1B, remaining cells were analyzed with the viSNE algorithm in Cytobank using the markers e-Cadherin and FITC to gate out the HCC827 labeled with eGFP. **B,** Phase-contrast microcopy of TAFs monocultures treated with 10 µM DON, 1 µM palbociclib or a combination of both showing that TAFs that have gained an activated CAFs-like morphology after treatment. **C,** Force-directed layout of TAFs mono- and cocultures individual samples, (**D)** TAFs monocultures showing proliferative (upper gate) and senescent (lower gate) TAFs and **(E)** TAFs size (FSC-A) in cocultures treated with 10 µM DON, 1 µM Palbociclib, a combination of both or without treatment.

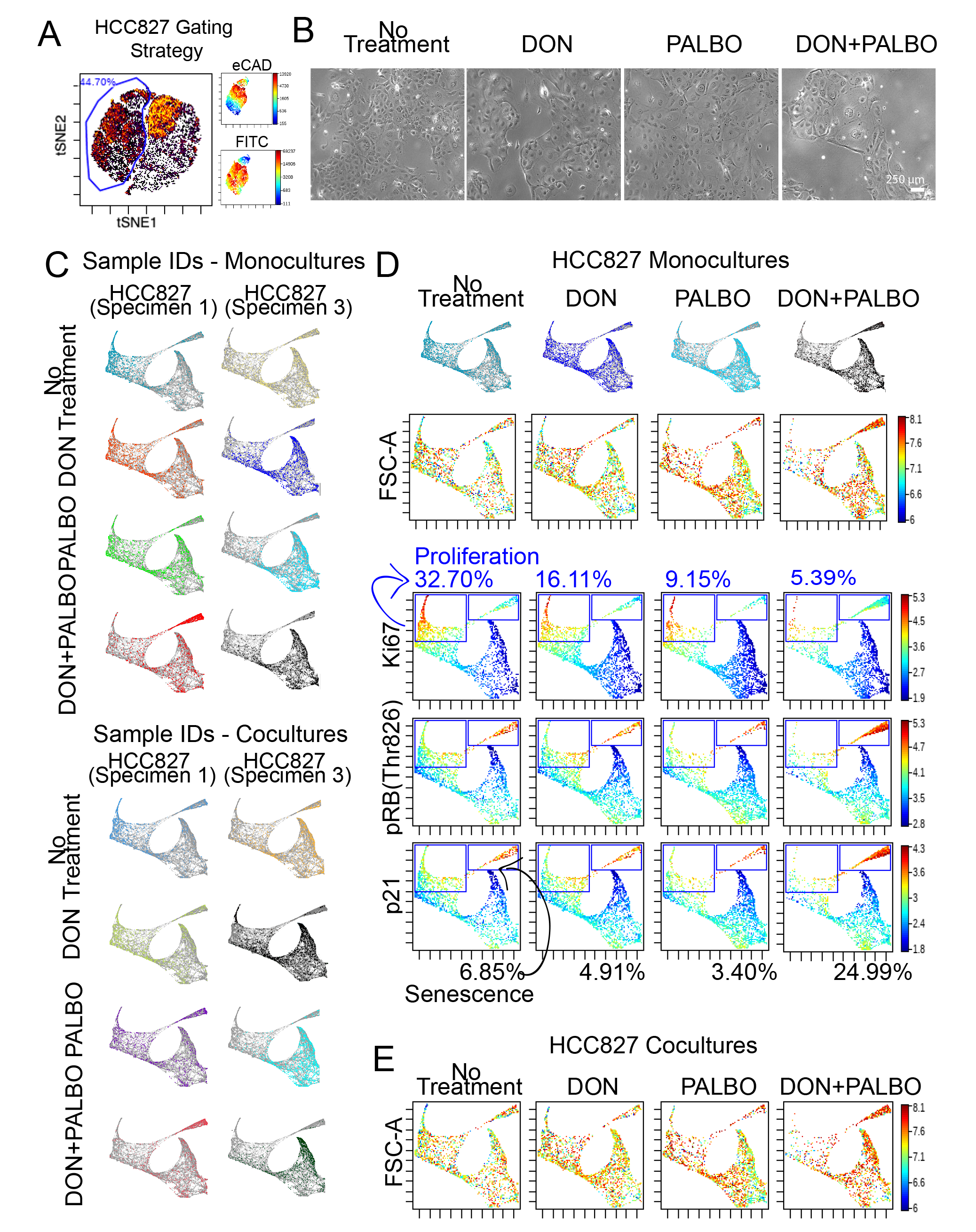

**Figure S5. Treatment with DON modulates Palbociclib efficacy in HCC827. (A)** Gating strategy for HCC827-eGFP. After removing debris, doublets and non-viable cells as shown in Figure S1B, remaining cells were analyzed with the viSNE algorithm in Cytobank using the markers e-Cadherin and FITC to gate the HCC827 labeled with eGFP. **(B)** Phase-contrast microcopy of the HCC827 monocultures treated with 10 µM DON, 1 µM palbociclib or a combination of both showing that HCC827 size increases after treatment. **(C)** Force-directed layout of HCC827 mono- and cocultures individual samples, **(D)** HCC827 monocultures showing proliferative (left gate) and senescent (right gate) HCC827s and **(E)** HCC827 size (FSC-A) in cocultures treated with 10 µM DON, 1 µM Palbociclib, a combination of both or without treatment.

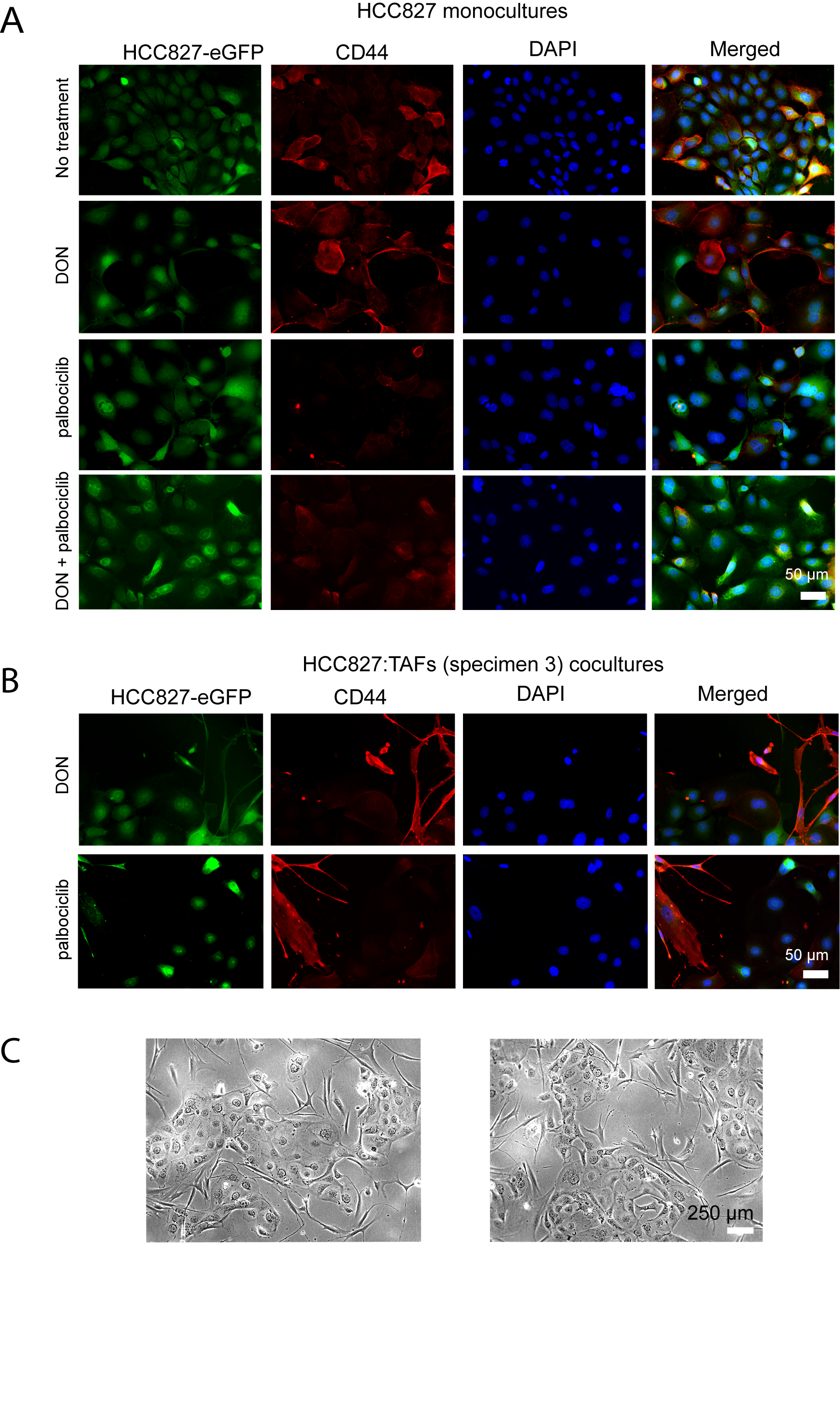

**Figure S6. Combination of DON and Palbociclib change TAFs coculture morphology. (A)** Immunofluorescence staining showing the mesenchymal marker CD44 in HCC827-eGFP monocultures with no treatment, DON, palbociclib, as well as in combination and **(B)** HCC827-eGFP:TAFs cocultures after treatment with DON and palbociclib alone. **(C)** Additional phase-contrast microcopy images of TAFs:HCC827 cocultures treated with DON and palbociclib showing a morphology similar to CAFs:HCC827 cocultures.

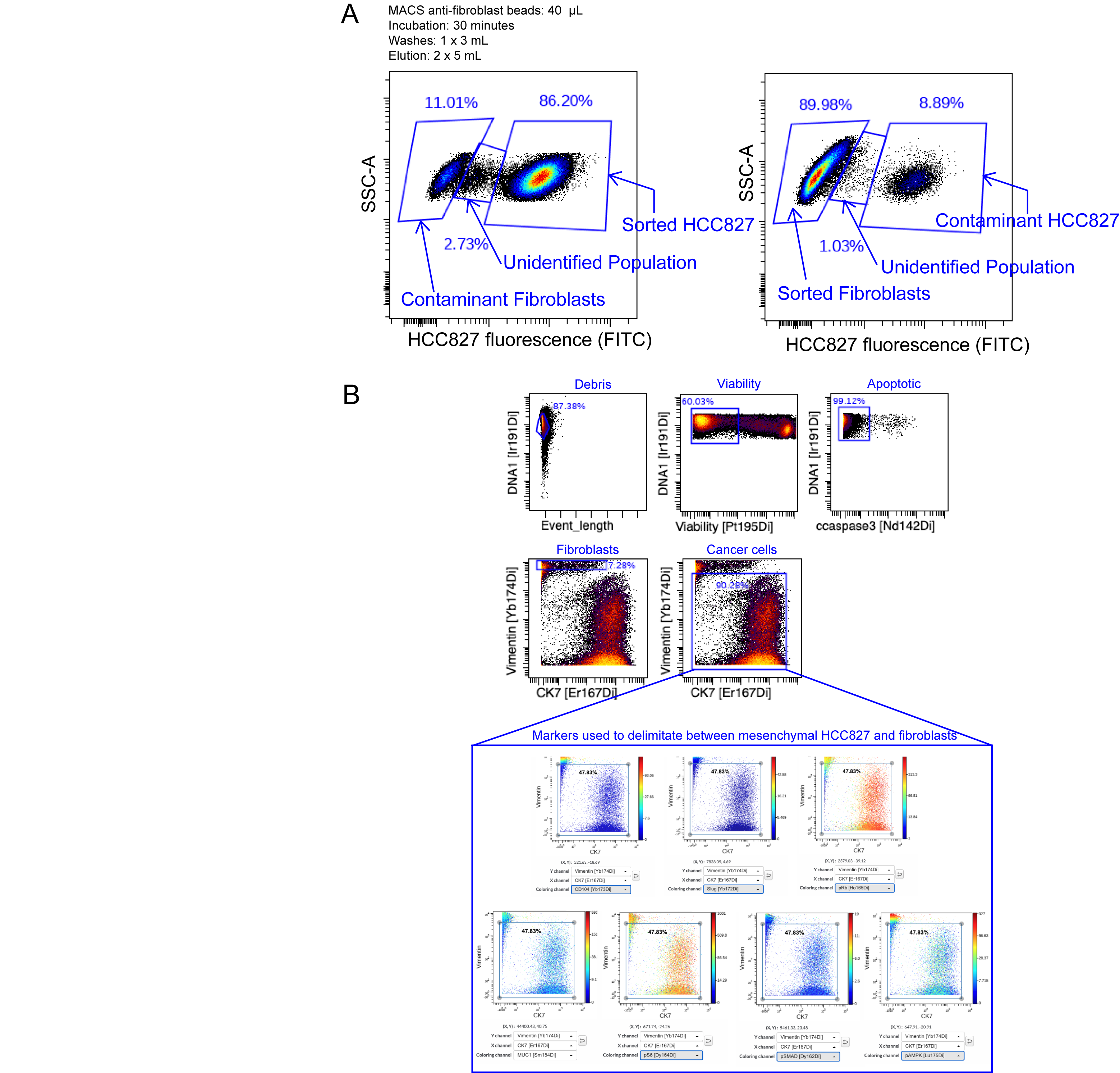

**Figure S7. MACS anti-fibroblast beads optimization and CyTOF gating strategy. (A)** MACS anti-fibroblast magnetic beads optimization protocol showing minimal contamination in each compartment after sorting HCC827-eGFP and primary LUAD fibroblasts cocultures. **(B)** After removing, debris, doublets and non-viable and apoptotic cells, we separated mesenchymal HCC827 from fibroblast cell population using known information about mesenchymal HCC827 from Karacosta et al. (Karacosta et al., 2019) including negative expression of CD104, Slug, and MUC1 as well as low signaling.
